# Variability in mitochondrial import, mitochondrial health and mtDNA copy number using Type II and Type V CRISPR effectors

**DOI:** 10.1101/2020.03.10.985606

**Authors:** Zuriñe Antón, Grace Mullally, Holly Ford, Marc W. van der Kamp, Mark D. Szczelkun, Jon D. Lane

## Abstract

Current methodologies for targeting the mitochondrial genome for basic research and/or therapeutic strategy development in mitochondrial diseases are restricted by practical limitations and technical inflexibility. The development of a functional molecular toolbox for CRISPR-mediated mitochondrial genome editing is therefore desirable, as this could enable precise targeting of mtDNA haplotypes using the precision and tuneability of CRISPR enzymes; however, published reports of “MitoCRISPR” systems have, to date, lacked reproducibility and independent corroboration. Here, we have explored the requirements for a functional MitoCRISPR system in human cells by engineering several versions of CRISPR nucleases, including the use of alternative mitochondrial protein targeting sequences and smaller paralogues, and the application of gRNA modifications that reportedly induce mitochondrial import. We demonstrate varied mitochondrial targeting efficiencies and influences on mitochondrial dynamics/function of different CRISPR nucleases, with *Lachnospiraceae* bacterium ND2006 (Lb) Cas12a being better targeted and tolerated than Cas9 variants. We also provide evidence of Cas9 gRNA association with mitochondria in HeLa cells and isolated yeast mitochondria, even in the absence of a targeting RNA aptamer. Finally, we present evidence linking mitochondrial-targeted LbCas12a/crRNA with increased mtDNA copy number dependent upon DNA binding and cleavage activity. We discuss reproducibility issues and the future steps necessary if MitoCRISPR is to be realised.

## INTRODUCTION

Mitochondria are crucial for the maintenance of cellular energy levels. In humans, all 37 genes contained within mitochondrial DNA (mtDNA) encode or aid biosynthesis of proteins involved in the OXPHOS pathway [1]. Despite mtDNA contributing only ~1% of mitochondrial proteins *in toto*—the remainder being encoded in the nucleus—four of the five OXPHOS complexes contain mtDNA-encoded subunits. Consequently, mutations in both nuclear and mitochondrial genomes can result in mitochondrial diseases with a range of clinical manifestations. Mitochondrial diseases are currently incurable, with a lack of understanding of the complexities of mitochondrial genetics and mitochondrial biology impeding identification of targets for the development of treatments [2]. Existing therapies mainly focus on symptom management through exercise and dietary supplements [3], [4]. Thus, the development of mitochondrial molecular genetics techniques that could be used therapeutically for mitochondrial diseases remains an important challenge.

Inherited disease-causing mtDNA mutations have been shown to be either homoplasmic, where near to 100% of mtDNA copies carry the mutation, or heteroplasmic, where the mutation is carried by a subset of the total mtDNA. In general, mutation load above ~70% is required to present a severe phenotype, although this threshold is disease specific. As a treatment to selectively degrade disease-causing mtDNA haplotypes, the field has developed targeted endonucleases which produce dsDNA breaks in the mutated mtDNA copies, which are then degraded by the mitochondria rather than being repaired [5, 6]. The remaining wild type mtDNA then replicates to re-establish mtDNA copy number. This “heteroplasmy purification” has been successfully demonstrated in cell culture and in mouse models using restriction endonucleases (MitoREs), zinc finger nucleases (MitoZFNs), TALENs (MitoTALENs), and homing endonucleases [7, 8]. However, although independently validated [9], these tools are impracticable and not widely adopted, because: (i) REs match few clinical mutations and cannot be readily re-engineered [10], and they additionally have high “off-target” cleavage rates [11, 12]; (ii) ZFNs require rounds of protein engineering/refinement; (iii) assembly of TALE parts is hindered by problematic cloning of DNA repeats; (iv) TALENs and ZFNs can have low import efficiency and can be mis-trafficked (e.g. ZFs have internal nuclear targeting sequences) [13–15]. To overcome some of these difficulties, we have investigated the use of a re-engineered flexible version of the CRISPR method for mitochondrial genome manipulation.

CRISPR systems have revolutionised molecular genetics by providing a quick and convenient method to efficiently target proteins to almost any short DNA sequence [16]. In this system, CRISPR guide RNAs (gRNAs) target effector nucleases to form a DNA/RNA hybrid (R-loop) at the target sequence, displacing the non-complimentary DNA strand and cleaving the two strands of DNA. Binding is assisted by a Protospacer Adjacent Motif (PAM) that is recognised by the CRISPR nuclease. There are several CRISPR-Cas system Types from different species, of which the Type II *Streptococcus pyogenes* (Spy) Cas9 and *Staphylococcus aureus* (SaCas9), and Type V *Lachnospiraceae bacterium* (Lb) Cas12a and *Acidaminococcus sp.* (As) Cas12a were tested in this study. Cas9 has been widely used for genome engineering. It comprises two nuclease domains, the RuvC-like and HNH domains, that generate blunt ended DNA double strand breaks (DSBs) [17, 18]. Cas12a, also referred to as Cpf1, contains a single RuvC endonuclease domain that cleaves the two DNA strands in turn, resulting in a staggered DSB with a 5′ overhang [19].

CRISPR-mediated manipulation of mitochondrial genomes requires both the CRISPR endonuclease and a gRNA (Cas9) or crRNA (Cas12a) need to be targeted to the mitochondria without compromising organelle viability (**Figure 1A**). To direct the import of non-native proteins, a naturally occurring mitochondria targeting sequence (MTS) is commonly cloned upstream of the protein of interest [20]. In contrast to the mitochondrial protein import pathway, the import of mitochondrial RNA is poorly understood [7]. Several studies have described methods to import RNA into mitochondria, either harnessing reportedly naturally occurring mitochondrial RNA import machineries or through non-natural means (reviewed in **Figure 1B,C**) [21–24]. Analogous to peptide MTSs, it has been proposed that short, structured RNA sequences (aptamers; e.g. from tRNAs, non-coding RNAs such as 5S ribosomal RNA, or the RNA component of RNase P) can be appended to an exogenous RNA for import e.g. [25, 26] (**Figure 1C**). However, there is no consensus on the mechanism(s) of RNA import, and the existence of a dedicated pathway for mitochondrial import of RNAs is debated [24, 25]. Consequently, the mitochondrial import of CRISPR gRNA/crRNA emerges as the critical challenge of developing MitoCRISPR.

**Figure 1.**
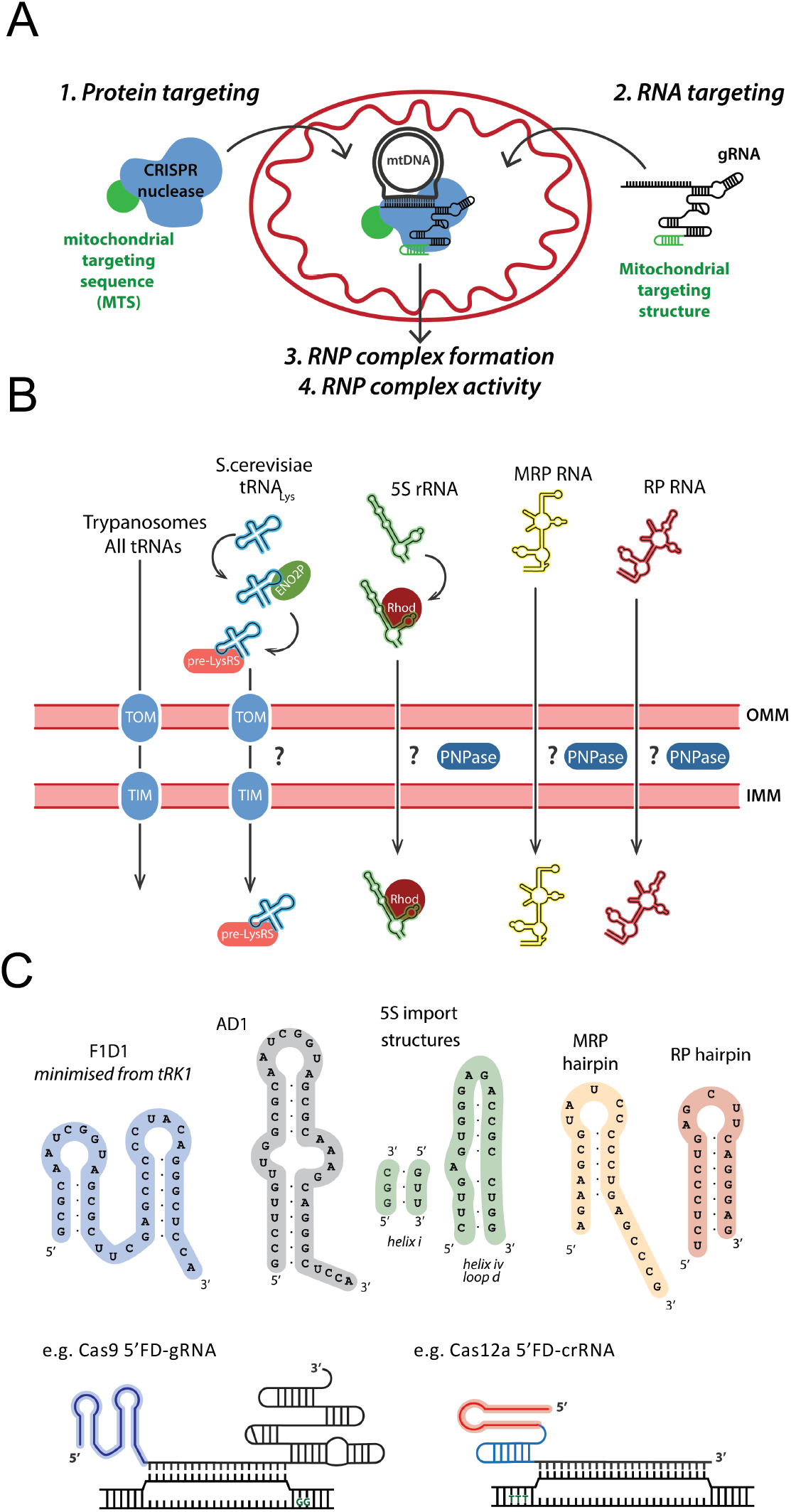
Mitochondrial targeting of CRISPR nucleases and RNAs. (A) The main steps required for the development of a working MitoCRISPR system: (i) targeted delivery of a CRISPR nuclease (1) and guide RNA (gRNA) (2) to the mitochondria; (ii) formation of a functional CRISPR ribonucleoprotein complex (RNP) inside the mitochondrial matrix (3); (iii) functional activity of the CRISPR RNP (4). (B) Natural RNA import mechanisms in Trypanosomes, *S. cerevisiae* and in mammals. (C) Minimised RNA import motifs that have been identified for tRK1, 5S rRNA, MRP RNA and RP RNA capable of directing RNA sequences of interest into mitochondria. Two examples are shown for Cas9 and Cas12a crRNAs modified with 5′FD and 5′RP respectively.

Steps towards development of CRISPR machinery for mtDNA have been described by others. Jo *et al* first introduced a standard nuclear targeted CRISPR/Cas9 system to edit mtDNA [27]. They unexpectedly reported that introducing nuclear-targeted Cas9 and unmodified gRNAs in HeLa cells resulted in mtDNA cleavage [27]. However, there have not been any published applications of this tool, and attempts to reproduce these data by us and by others [28] have not been successful. The second report was published by Orishchenko *et al* using a FLAG-Cas9 targeted to fragmented mitochondria of viral-producing cells by appending a COX8A MTS [29, 30]. The paper showed only one cell in which anti-FLAG colocalised with very fragmented mitochondria and did not describe any experiments to address mitochondrial gRNA localisation. As with the work of Jo *et al*, these data have not yet been independently corroborated. A recent publication on MitoCRISPR attempted to add mitochondrial-targeting structures to the gRNA for the first time [25]. Loutre *et al* showed a decrease in mtDNA copy number with COX8A-Cas9 and gRNAs modified to contain the targeting sequences from 5S RNAs. Two gRNAs were required to produce a reduction of mtDNA copy number; a single target gRNAs had no significant influence on mtDNA content [25]. The work was carried out in a heteroplasmic cell line, but the authors did not observe a shift in heteroplasmy.

The explosion in gene editing research more widely has been driven by the simplicity, reproducibility and adaptability of CRISPR-Cas systems. Almost immediately after the first descriptions of CRISPR Cas9, multiple labs demonstrated reliable nuclear gene editing by the facile recoding of a gRNA/crRNA spacer sequence. This has not so far been the case with MitoCRISPR, and we suggest that possible damage to mitochondria during cell fractionation and mitochondrial isolation, and/or failing mitochondrial proteostasis due to Cas9-expression, might explain some or all of the reports of MitoCRISPR-type activity in previous publications. It is critical that MitoCRISPR is fully characterised (mtDNA copy number reduction alone is not sufficient) and that there is clear evidence of reproducibility, validated by independent labs.

By designing and re-engineering several versions of CRISPR nucleases and gRNAs, including testing alternative MTSs, smaller paralogues, and the addition of mitochondrial targeting RNA aptamers, we have assessed: (i) CRISPR/Cas mitochondrial targeting and influence on mitochondrial dynamics/function; and (ii) DNA cleavage activity in a model mitochondrial disease cell line (MELAS cybrids). We explore the requirements for targeting of Cas9 gRNA to mitochondria, including the influence of appending RNA aptamers. Our study shows that SpyCas9 can be localised to mitochondria, but with low efficiency and causing severe mitochondrial network damage. However, LbCas12a was found to be efficiently expressed and targeted to the mitochondrial matrix via the TOM/TIM pathway in WT or MELAS cybrids. Finally, we present our attempts to demonstrate an effect on mtDNA of a mitochondrial-targeted LbCas12a/gRNA combination. While we observed increases in mtDNA copy number in MELAS disease model cybrid, dependent upon DNA binding and cleavage activity, this had poor reproducibility across independent validations and stable/prolonged Cas12a expression had deleterious effects on respiration. Our data emphasise the importance of CRISPR enzyme and mitochondrial targeting strategy selection, and further highlight how manipulation of the mitochondrial genome can have unpredictable outcomes in living cells.

## MATERIAL AND METHODS

### Reagents and antibodies

Unless stated otherwise, all reagents were from Sigma. Stock solutions of CCCP (carbonyl cyanide m-chloro phenyl hydrazone; C2759; 10 mM), DAPI (4’, 6-diamidino-2-phenylindole; D9542; 1 mg/mL), MB-10 (MCule; P-579567132), MB-12 (PHR1300-500MG), cycloheximide (TOKU-E, C084) and proteinase K (Thermo, EO0491; 10 mg/mL) were stored at −20°C. The following primary antibodies were used: anti-GAPDH (G8796); anti-GFP for immunoblotting (Covance, MMS-118R); anti-GFP for immunoEM (Rockland, 600-401-215); anti-HSP60 (H4149); anti-Tom20 (BD Biosciences, 612278); anti-tubulin (T5168); polyclonal anti-LbCas12a (GeneTex, GTX133301); monoclonal anti-LbCas12a (SAB4200777). HRP-tagged secondary antibodies were from Jackson ImmunoResearch; fluorescent Alexa-tagged antibodies were from Molecular Probes. VOA: 1 mM valinomycin, 10 mM oligomycin, 8 mM antimycin A.

### Cell-lines and cell culture

hTERT-immortilised human Retinal Pigment Epithelial (RPE1) cells were maintained in high-glucose DMEM medium (D5796) supplemented with 10% FBS (F7524) at 37°C in 5% CO_2_. Wild type ACH cytoplasmic hybrid cells or 94αT3 MELAS cybrids carrying m.A3243G mtDNA mutation (a kind gift from Dr José Sanchez-Alcázar, Seville [31]) were maintained in high-glucose DMEM medium supplemented with 5% (v/v) FBS, 1 mM sodium pyruvate, 1% (v/v) penicillin/streptomycin (P4333) and 50 μg/mL uridine at 37°C in 5% CO_2_.

### Plasmids and transfection

Transient transfections were performed with Lipofectamine 2000 Reagent (Thermo Scientific, 11668027) and OptiMEM reduced serum medium (Gibco) according to the manufacturer’s instructions. The efficacy of cell transfection was checked using fluorescence microscopy. Plasmid sequences are available on request: pD1301-AD (DNA2.0), for expression of NLS-SpyCas9-GFP or COX8a-SpyCas9-GFP and unmodified Cas9 gRNA template generation; pEX-A2 (Eurofins), for 5′RP/5′FD gRNA template generation; pSP1(Siksnys lab [32]), used as a plasmid cleavage assays substrate plasmid; pEGFP (Clontech), for expression of COX8a/ATG4D-(SaCas9/LbCas12a/AsCas12a)-GFP or LbCas12a-GFP; Su9-GFP (Addgene, #23214 [33]), for expression of Su9-(SaCas9/LbCas12a/AsCas12a/PI)-GFP and Su9-(mutant/dLbCas12a)-GFP; pLVX-puro (Clontech), used for stable expression of Su9-(LbCas12a)-GFP; PLKO.1 (Addgene, #10878 [34]), for expression of Cas12a gRNAs; SaCas9, LbCas12a, AsCas12a, dLbCas12a (Addgene, #61592 [35], #69988 [36], #69982 [36], #104563 [37]), for protein template generation.

### Viruses, transduction and stable cell-lines

Lentiviruses were generated in HEK293T cells by transient transfection using PEI reagent. 27 μg of the plasmid of interest was transfected together with 20.4 μg of the packing plasmid pAX2 and 6.8 μg of the envelope plasmid pMGD2. Viruses were harvested 48 h after transfection. Media were collected and centrifuged 1,500 x *g* for 5 min and filtered with a 0.45 μm filter to remove cells and debris. Viruses were concentrated using Lenti-X Concentrator (Clontech). One volume of Lenti-X Concentrator was combined with three volumes of clarified supernatant. The mixture was incubated 1 h at 4°C, then centrifuged at 1,500 x *g* for 45 min at 4°C and the pellet resuspended in N2B27 media or DMEM media. For viral transduction, cybrids were plated in 6cm dishes and transduced with the corresponding lentiviruses (7.5 μl/mL) in the presence of 8 μg/mL polybrene to increase transduction efficiency. After two days, GFP-positive cells were selected by fluorescent activated cell sorting (FACS). Stable line generation was checked using fluorescence microscopy.

### Immunoblotting

Cells grown on 6-well plates were initially washed with ice-cold PBS, then lysed with 100-200 μL/well of ice-cold radioimmunoprecipitation assay (RIPA) buffer consisting of 50 mM Tris HCl (pH7.4), 1% Triton-X-100 (9002-93-1), 0.5% sodium deoxycholate (D6750), 150 mM NaCl (S9888), 0.1% sodium dodecyl sulphate (SDS, 436143) supplemented with one tablet of protease inhibitor per 10mL of RIPA buffer. The homogenates were incubated on ice for 15 min, then cleared by centrifugation at 12,000 x *g* for 15 min at 4°C. Supernatants were collected as soluble fractions. Proteins were transferred to nitrocellulose membranes (Biolabs, 1620115). Membranes were then incubated with primary antibody diluted in 5% milk or 2.5% BSA in Triton X-100-TBS buffer (T-TBS) overnight. Primary and secondary antibodies used are listed above. Membranes were then washed three times prior to incubation with ECL Chemiluminiscence reagents (Geneflow, K1-0170), and band intensities were detected in films (GE Healtchare, 28906837) using a film developer.

### Subcellular fractionation and proteinase K treatment

Cybrid cells were plated on 10 cm dishes and transfected with the Su9LbCas12a-GFP construct using Lipofectamine2000 for 48 h. Cells were trypsinized, washed in PBS and resuspended in mitochondrial import buffer (50 mM HEPES, pH 7.1; 0.6 M sorbitol; 2 mM KH_2_PO_4_; 50 mM KCl; 10 mM MgCl_2_; protease inhibitors) [38]. Cells were lysed on ice using a ball bearing cell homogeniser (twenty passes at 10 μm clearance; Isobiotech, Heidelberg, Germany) then centrifuged at 800*g* for 5 min to remove the heavy nuclear pellet. The postnuclear supernatant was then centrifuged for 15 min at 10000 x *g*, and the resulting mitochondria-containing pellet was washed with 140 μL import buffer and incubated in the absence or presence of 0.1mg/mL proteinase K for 15 min on ice. The reaction was stopped by the addition of phenylmethylsulfonyl fluoride (PMSF) to 1 mM.

### Immunofluorescence and microscopy

Cells were seeded on coverslips. Cells were washed twice with PBS and incubated with 4% formaldehyde for 15 min or −20°C methanol for 5 min. Cells were then incubated for 30 min with primary antibody (listed above) in PBS. Cells were washed three times with PBS and incubated with the secondary antibodies (listed above) and counterstained with DAPI (Life Technologies, D121490, 100ng/mL) for 10 min. Cells were then washed again with PBS and mounted in Mowiol. Fixed-cell images were obtained using an Olympus IX-71 inverted microscope (60x Uplan Fluorite objective; 0.65-1.25NA, oil immersion lens) fitted with a CoolSNAP HQ CCD camera (Photometrics, AZ) driven by MetaMorph software (Molecular Devices). Confocal microscopy was carried out using a Leica SP5-AOBS confocal laser scanning microscope (63x oil immersion objective, 1.4NA; or 100x oil immersion objective, 1.4NA) attached to a Leica DM I6000 inverted epifluorescence microscope. Laser lines were: 100 mW Argon (for 458, 488, 514 nm excitation); 2 mW Orange HeNe (594 nm); and 50 mW diode laser (405nm). The microscope was run using Leica LAS AF software (Leica, Germany). Image analysis was performed using Fiji (National Institutes of Health, Bethesda, USA).

### Analysis of DNA or cDNA levels by quantitative real-time polymerase chain reaction (qRT-PCR)

Cybrid cells were plated on 6 cm dishes (DNA) or 6-well plates (cDNA). Cells were transiently transfected for 2 days. For DNA extraction, cells were then washed with ice-cold PBS and lysed with 200 μL lysis buffer (1% SDS, 10 mM EDTA and 50 mM Tris pH 8.1) and incubated on ice for 10 min. For the purification of DNA, the phenol-chloroform purification was used. Finally, the DNA pellet was resuspended in a final volume of 15 μL and diluted for qRT-PCR analysis with specific primers. For RNA extraction, after the corresponding transfection, cells were washed with PBS and then cells were lysed in 350 μL RLT buffer (Qiagen). Total RNA was extracted through columns using RNeasy kit (Qiagen, 74104) following manufacturer’s instructions and genomic DNA was digested using DNaseI (Qiagen). RNA samples were reverse transcribed using High-Capacity RNA-to-cDNA™ Kit (Thermo Scientific, 4387406), according to manufacturer’s protocol. The DNA or cDNA samples were amplified using SYBR Green (Life Technologies). The reaction was carried out using StepOnePlus System (Applied Biosystems) and the following conditions were selected: after an initial denaturation at 95°C for 10 min, 40 cycles with 95°C for 15 s (denaturation), 60°C for 30 s (annealing) and 60°C for 30 s (elongation). For the analysis, DNA or mRNA levels were estimated using the ΔΔCt method normalising data to GAPDH levels [39].

### Seahorse bioenergetics

Cybrid cells were plated according to manufacturer’s instructions on 8-well Seahorse XFp plates (Agilent, 103010-100). The day of the assay, culture media was replaced with Seahorse XF base medium (Agilent) supplemented with 1 mM sodium pyruvate, 2 mM glutamine, and 10 mM glucose (pH 7.4) for 1 h at 37°C. The Mito Stress Test Kit (Agilent) was prepared according to the manufacturer’s instructions: oligomycin (1 μM); carbonilcyanide ptriflouromethoxyphenylhydrazone (FCCP; 1 μM); rotenone/antimycin A (0.5 μM). After analysis, cells were lysed in 20 μl of RIPA buffer and protein levels were quantified by Nanodrop (A280) to normalise the data.

### *In vitro* transcription of 33P-UTP labelled RNA

*In vitro* transcription was performed using the T7 HiScribe High Yield RNA synthesis kit. The reaction was prepared to a final volume of 20 μL, containing a final concentration of 1X reaction buffer, 1 mM ATP, GTP and CTP. UTP was added to a final concentration of 4 μM with 0.25 μM α^33^P UTP, 0.5 MBq/reaction (Hartman Analytic, SRF 210). To this, 0.25 μg template DNA and 1 μL T7 RNA pol mix was added. The assembly was incubated at 37°C for 10 min, treated with DNase and purified with MicroBioSpin (BioRad) columns.

### Denaturing urea-PAGE

RNA was separated on a denaturing urea-PAGE gel containing 10% (w/v) acrylamide, 1 X Tris-Borate EDTA (TBE) and 50% (w/v) urea. The gel mix was filter sterilised with a 0.2 μm nitrocellulose membrane. Gels were set using the BioRad Mini-PROTEAN 3 system and pre-run in 1 X TBE at 200 V for 45 min prior to loading samples. RNA samples were heated to 90 °C with 1X RNA Loading Dye for 10 min and migrated in 1 X TBE at 200 V. Gels were stained with 1 mL 1/1,000 SYBR Gold Nucleic Acid Gel Stain (ThermoFisher Scientific) and imaged with the GelDoc™ XR+ system (BioRad).

### CRISPR (crRNA/Cas12a) complex assembly and cleavage assay

250 nM *Lachnospiraceae bacterium* Cas12a protein was mixed with 250 nM crRNA in SB buffer (10 mM Tris pH 7.5, 100 mM NaCl, 1 mM EDTA, 0.1 mM DTT, 5 μg/mL BSA) and incubated at 37°C for 1 h. For each cleavage reaction, 3 mM plasmid substrate (pSP1) was assembled in RB buffer (10 mM Tris pH 7.5, 10 mM Mg2Cl, 1 mM EDTA, 0.1 mM DTT, 5 μg/mL BSA) at 37°C and preheated for 5 min. The reaction was started by addition of 50 nM assembled Cas12a CRISPR complex which was incubated for the time period specified. The reaction was quenched by adding 1/3 volume of 80°C 3 X STEB (0.05 M Tris-HCl pH 8.0, 0.05 M EDTA, 20% (w/v) sucrose, 0.0125% (w/v) bromophenol blue, 0.0125% (w/v) xylene cyanol) and incubating at 80°C for 5 min. Samples were separated by electrophoresis on a 1.5% (w/v) agarose gel stained with ethidium bromide at 20 V overnight (16 h) and visualised by UV irradiation.

### Isolation of yeast mitochondria

Yeast (YPH499) were grown in 400 mL YPG with Pen/Strep at 24°C, 150 rpm for ~48-72 hours. When OD600 = 7.5, cells were sub-cultured into 2 × 1 L YPG pH 5.2 supplemented with Pen/Strep to OD600 = 0.5, then grown overnight at 19°C, 120 rpm. When OD600 = 1.2-1.8, cells were pelleted (4,000 × *g*, 10 min, RT) and pellets re-suspended in dH_2_O. Mitochondria were isolated through differential centrifugation after cell wall was reduced with DTT as described in [40].The final mitochondrial pellet was resuspended in 250 mM sucrose and 10 mM MOPS, pH 7.2. Mitochondrial protein was quantified by BCA assay, using BSA as standard. 1mg aliquots were snap frozen and stored at −80°C.

### Import of radiolabelled RNA into isolated yeast mitochondria

Isolated yeast mitochondria were thawed on ice and diluted in ice cold yeast import buffer (50 mM sucrose, 80 mM KCl, 5 mM MgCl_2_, 10 mM KH_2_PO_4_/K_2_HPO_4_, 10 mM MOPS, pH 7.2) with 2 mM ATP and 2 mM NADH added just before use. 100 μg mitochondria were diluted into 100 μL import buffer per condition. The mitochondrial suspension was preheated to 25°C in a thermo-mixer with gentle shaking and import started by addition of ^33^P-labelled RNA, gentle vortex and incubation at 25°C. Import was stopped after 5 min with VOA. The sample was split into two Eppendorf tubes (50 μL each) and one tube incubated with 5 μg RNase A for 15min at 25°C. RNase was stopped with 1.5 μL RNase inhibitor and incubated for 10 min at 25 °C, then washed twice with 250 μL yeast import buffer with 1 μL RNase inhibitor. Samples were centrifuged at 17,000 x *g*, 10 mins, and washed pellets either run on a denaturing urea-PAGE gel directly, or the RNA extracted with TRIzol reagent. For TRIzol extraction of RNA: each pellet was resuspended in 250 μL TRIzol reagent and tubes were incubated at RT for 5 min, 50 μL chloroform was added and tubes incubated for a further 5 min at RT. Samples were centrigued for 15 min at 12,000 x *g*, 4 °C, and the upper aqueous phase transferred to a new tube with 125 μL isopropanol with 0.05 μg/μL glycogen and incubated for 10 min at RT. Precipitated RNA was pelleted by centrifugation at 12,000 x *g* for 10 min, 4°C, and pelleted RNA was washed twice with 250 μL 75% (v/v) EtOH (7,500 *g*, 5 min, 4°C), air dried and resuspended in 10 μL 2X formamide RNA loading buffer. For direct separation by denaturing-PAGE, pellets were resuspended in 5 μL yeast import buffer and 5 μL 2X formamide RNA loading buffer. RNA in formamide RNA loading buffer was heated to 100°C for 10 min before separation by denaturing urea-PAGE. The gel was fixed in gel fixing solution (10% (v/v) MetOH, 10% (v/v) acetic acid) for 1 h, then 30 min gel fixing solution with 5% (v/v) glycerol. The gel was dried with a pre-programmed schedule (up to 80°C, over 2 h) on a BioRad model 583 gel dryer, and then exposed on a phosphorimager screen (Fujifilm). Phosphor screen was imaged with a Typhoon-Trio scanner (GE Healthcare).

### Cellular fractionation and isolation of RNA for Northern blotting

HeLa cells were seeded to be 90% confluent on the day of the fractionation. Following aspiration of culture media, cells were washed two times with PBS. Cells were detached with trypsin and transferred into a 14 mL falcon tube and centrifuged at 600 *g* at 4 °C for 10 min. The cell pellet was resuspended in 1 mL icecold isolation buffer (0.44 M sorbitol, 40 mM EDTA, 10 mM HEPES-NaOH (pH 6.7), 0.1 % (w/v) SDS). Cells were homogenized using a cell homogenizer with an 8 μm spacing ball bearing, according to the manufacturer’s instructions. 100 μL of whole cell extract was kept for later RNA extraction. The remaining homogenate was centrifuged at 1,500 x *g* for 5 min at 4°C to pellet nuclei. The supernatant was collected and centrifuged at 15,000 x *g* for 20 min at 4°C to pellet mitochondria. The cytosolic fraction was kept on ice and the mitochondrial pellet was washed with isolation buffer and resuspended in 300 μL isolation buffer prior to disruption of the outer mitochondrial membrane with 60 ng/μL digitonin for 7 min. at RT. Digitonin was diluted by a wash with 700 μL isolation buffer, and after a final wash the mitochondrial pellet, nuclear pellet, cytoplasmic fraction and whole cell sample were resuspended in 500 μL TRIzol and stored at −80 °C prior to RNA extraction as per the manufacturer’s instructions.

### Northern blotting

RNAs were electro-transferred from urea-PAGE gels to Hybond XL nylon membrane (GE Healthcare) at 12 V over 12 h in 0.5X TBE for Northern hybridisation. RNAs were fixed onto nylon membrane by UV irradiation at 1,500 x 100 μJ/cm^2^. Membranes were prehybridised in Buffer NB1 (0.1% (w/v) SDS, 0.9 M NaCl, 0.09 M sodium citrate, 0.3 mM EDTA, 0.2% (w/v) BSA), 0.2% (w/v) Ficoll 400, 0.2% (w/v) polyvinylpyrrolidone). Hybridisation of 8 × 7 cm membranes in 50 mL falcons used 2.9 mL Buffer NB1. Membranes were hybridised in one volume of Buffer NB1 containing one volume ([probe]f: 2 ng/mL) 5’-^32^P-labelled oligonucleotide probe by incubation overnight at 5°C. Probes (**Table 2**) of known cytoplasmic and mitochondrial RNAs were used as positive controls for cellular compartments, and designed hybridisation probes against gRNAs and mitochondrial targeted RNA aptamers determined transfected RNA cellular localisation. Following hybridisation, membranes were washed three times for 10 min in Buffer NB2 (0.1% w/v) SDS, 0.3 M NaCl, 0.03 M sodium citrate, 0.1 mM EDTA) at room temperature and exposed on a phosphorimager screen which was imaged with a Typhoon-Trio scanner (GE Healthcare).

### Labelling hybridisation probes

Dephosphorylated DNA oligonucleotides were labelled with bacteriophage T4 polynucleotide kinase by incubation for 1 h at 37°C with a 5-fold molar excess of 10 mCi/mL γ-^32^P ATP (Hartman Analytic, SRP 301) as per the manufacturers’ instructions. The reaction was terminated by addition of EDTA (final concentration 20 mM), and probes were purified from unincorporated nucleotide with BioRad Micro Bio Spin P-6 columns.

### Statistics

Graphical results were analysed with GraphPad Prism 7 (GraftPad Software, San Diego, CA), using an unpaired Student’s t-test or one-way ANOVA, as indicated. *p<0.05, **p<0.01 and ***p<0.001. Results are expressed as mean ± SEM or mean ± SD, as indicated.

## RESULTS

### Design of a MitoCRISPR system

The development of a reliable MitoCRISPR system requires (**Figure 1A**): (1) the targeted delivery of a CRISPR nuclease to the mitochondria by fusion to a peptide mitochondrial targeting sequence (MTS); (2) mitochondrial targeting of a CRISPR gRNA, possibly through fusion to a mitochondrial targeting domain; (3) formation of a functional CRISPR ribonucleoprotein (RNP) complex inside the mitochondrial matrix; and (4) functional nuclease activity of the CRISPR RNP complex. There are multiple ways in which a CRISPR endonuclease protein could be delivered into the cell for subsequent import into mitochondria. In the context of this project, CRISPR proteins were delivered as DNA, either stably expressed by incorporation into the genome or transiently transfected from a plasmid. We tested the previously used SpyCas9, less commonly used SaCas9, as well as Type V LbCas12a and AsCas12a. The addition of an MTS was based on extensive work targeting other proteins for mitochondrial import, and here we tested three different well-characterised signal sequences. The requirement for protein unfolding prior to mitochondrial import would appear at this stage to rule out ribonucleoprotein transfection as a potential alternative strategy (see discussion below).

Several functional non-coding RNAs (ncRNAs) are thought to be imported into the mitochondria, including tRNAs, the 5S RNA, MRP RNA, RNaseP RNA and mitochondrial micro RNAs (MitomiRs) (**Figure 1B**). Although whole transcriptome analyses of mitochondria and mitoplasts have confirmed the presence of these RNAs [41, 42], the mechanisms of import and impact of these ncRNAs are not fully explored. Mitochondrial RNA import pathways differ from one organism to another. Trypanosomes import all their tRNAs through TOM/TIM, whereas *S. cerevisiae* import just one, tRNALys. Import of tRNALys involves the protein factors enolase (ENO2P) and pre-LysRS, the pre-cursor of the mitochondrial lysyl-tRNA, acting in concert with a functional protein import machinery [43, 44]. In mammalian cells, mitochondrial import of 5S ribosomal RNA (rRNA), mitochondrial RNase P RNA (MRP), and RNase P RNA (RP) occurs by unknown mechanisms, although evidence implicates factors including Rhodanese (Rhod) in the case of the 5S rRNA [45–47], and possibly PNPase during 5S, MRP and RP RNA import [48–50] (**Figure 1B**). Despite conflicting evidence on the existence and mechanisms of mammalian RNA import, several groups have proposed minimal structural motifs that are sufficient to direct import (**Figure 1C**). The F1D1 RNA, derived from tRK1, is the structure found to have the highest import efficiency. A related control sequence, the AD RNA, does not have the same effect [23]. Moreover, truncating RNase RP and MRP RNAs has allowed identification of similar approximately 20nt RNA hairpins in both RP and MRP RNA which direct mitochondrial import. Attaching these hairpins into non-imported RNAs can target RNAs for import, as shown with GAPDH [51]. In the context of this study, either the FD, AD or RP motifs were used for the import of Cas9 gRNA or Cas12a crRNAs into mitochondria by attachment at the 5′ends (**Figure 1C, Table 1**).

### Targeting of CRISPR/Cas9 to mitochondria

#### SpyCas9 gRNAs co-fractionate with mitochondria in HeLa cells and isolated yeast mitochondria both with and without a targeting aptamer

The subcellular localisation of modified Cas9 gRNAs was determined using a Northern blot method (**Figure 2A**). A cytoplasmic RNA (tRNALys) and a mitochondrial RNA (tRNALeu) were used as control markers. Probes designed against the corresponding transfected gRNA are given in **Table 2**. Cells were transfected with unmodified gRNA, gRNA with 5′RP, and gRNA with 5′FD, and the hybridisation signal was determined using a probe for the hairpin region of the gRNA (see **Figure S1**). Strong signals for all three gRNAs were observed in the nuclear fraction and, to a lesser extent, the mitochondrial fraction (**Figure 2B**). The 5′FD gRNA produced the weakest signal in the mitochondrial fraction, contrasting with a stronger signal for the 5’RP and the unmodified gRNA. Note that the cytoplasmic tRNA probe also gave a signal for the mitochondrial fraction, and both cytoplasmic and mitochondrial tRNA probes gave strong nuclear fraction signals (**Figure 2B**).

**Figure 2.**
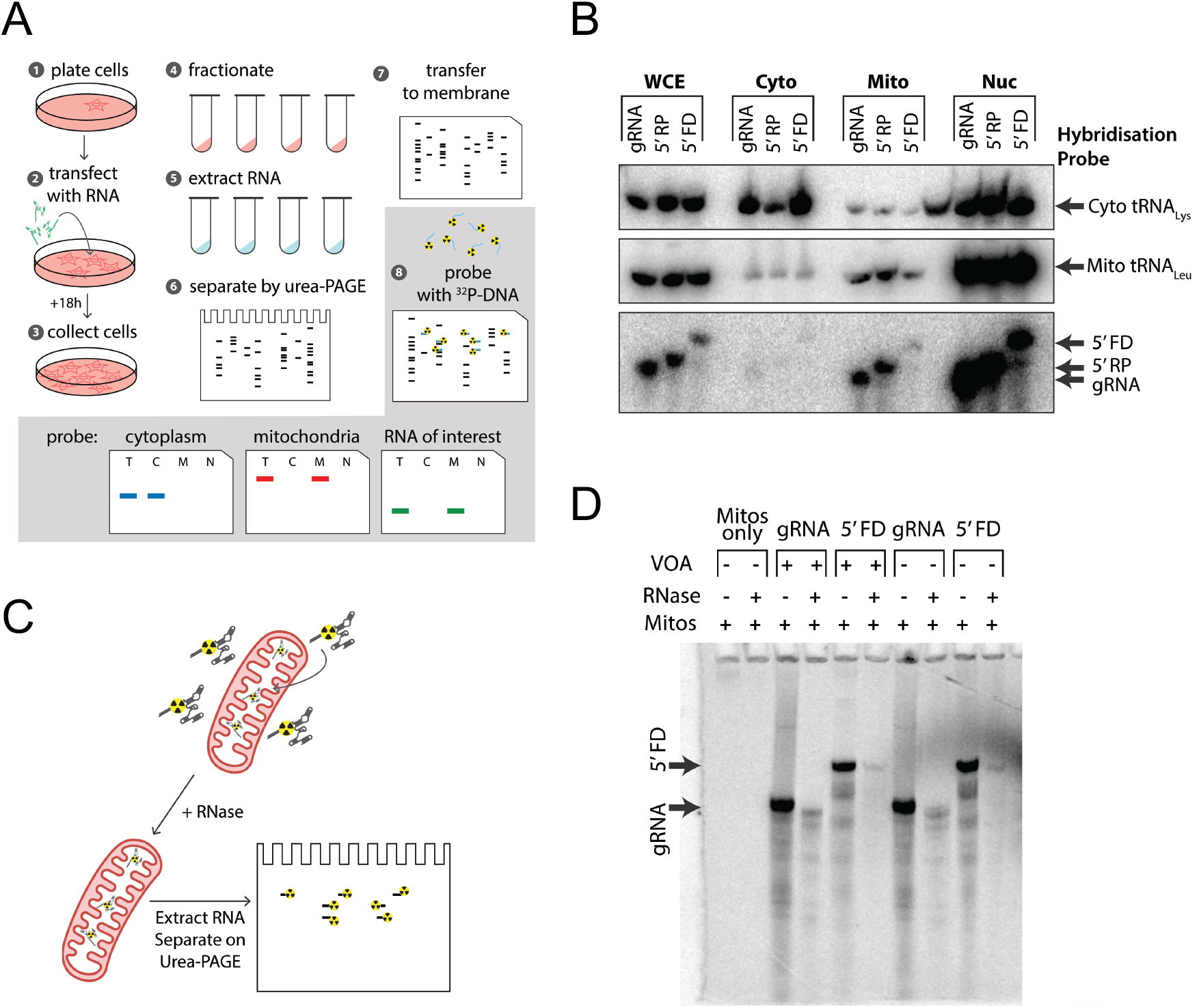
Import of Cas9 gRNAs into mitochondria in HeLa cells or isolated yeast mitochondria. (A) Northern blotting workflow: cells are plated in a 10 cm dish and transfected with RNA when 70-80% confluent. After 18 hours, cells are harvested, homogenised and fractionated. RNA is extracted from fractions and separated by urea-PAGE. Separated RNAs are transferred to nitrocellulose membrane and probed with ^32^P-DNA. (B) Northern blotting for cells transfected with either unmodified gRNA, or gRNA modified with 5′RP or 5′FD; Lanes were loaded according to fractionation: whole cell extract (WCE); cytoplasmic (Cyto); Mitochondrial (Mito); nuclear (Nuc). The three blots represent different northern probes: cytoplasmic tRNA_Lys_ (*Top*); mitochondrial tRNA_Lys_ (*Middle*); gRNA (*Bottom*). The different mobilities of the gRNA are due to the differences in size when RP or FD aptamers are present. (C) Assessing RNA import in isolated mitochondria: RNA import can be measured by directly importing radiolabelled RNAs. (D) Import of radiolabelled RNA into isolated yeast mitochondria. TRIzol extracted RNA was separated on urea-PAGE. Following separation, gels were fixed, dried and exposed to a phosphor screen.

To provide a sensitive method for following RNA import into isolated mitochondria, gRNAs were *in vitro* transcribed with ^33^P-UTP. Unmodified gRNA and a modified gRNA were incubated with mitochondria with or without a membrane potential (±VOA). Following a 10-minute incubation at 25°C, pelleted mitochondria were treated with RNase to degrade non-imported RNA. RNA was then extracted using TRIzol, and samples were separated on urea-PAGE (**Figure 2C, D**). **Figure 2D** shows that a portion of both unmodified and 5′FD modified gRNAs was protected from RNase-mediated digestion, presumably by the mitochondrial membrane(s). Partial gRNA protection was observed with VOA treatment, which could be due to RNA protection by membrane association rather than import. The protected gRNA bands migrated more rapidly than corresponding bands before treatment; this contrasted with the FD gRNA that appeared as a single faint band of the same size (**Figure 2D**).

In both assays, gRNA appear to be associated with mitochondria regardless of whether an RNA import aptamer was attached. This could provide support for the possibility that MitoCRISPR can be achieved without targeted import of the RNA. However, we note that these assays do not report on RNA import to the matrix, and in both cases association with the outer membrane or partial insertion could give similar results.

#### SpyCas9 is localised to mitochondria with low efficiency and causes mitochondrial damage

**Figure 3A** shows the two SpyCas9 proteins used in this project which we expressed from human codon optimised genes: nuclear targeted SpyCas9 (NLS-SpyCas9-GFP); mitochondria targeted SpyCas9 (MTS-SpyCas9-GFP); and a control protein, mitochondria targeted GFP (MTS-GFP). We first tested the N-terminal MTS from human cytochrome c oxidase subunit viii (COX8A), one of the most widely used signal peptides comprising a 29-residue helix. Different transfection conditions, cell fixation methods, and mitochondrial markers were used to assess the targeting efficiency and influence of mitochondria targeted Cas9 in human hTERT RPE1 cells (**Figure 3B-E**).

**Figure 3.**
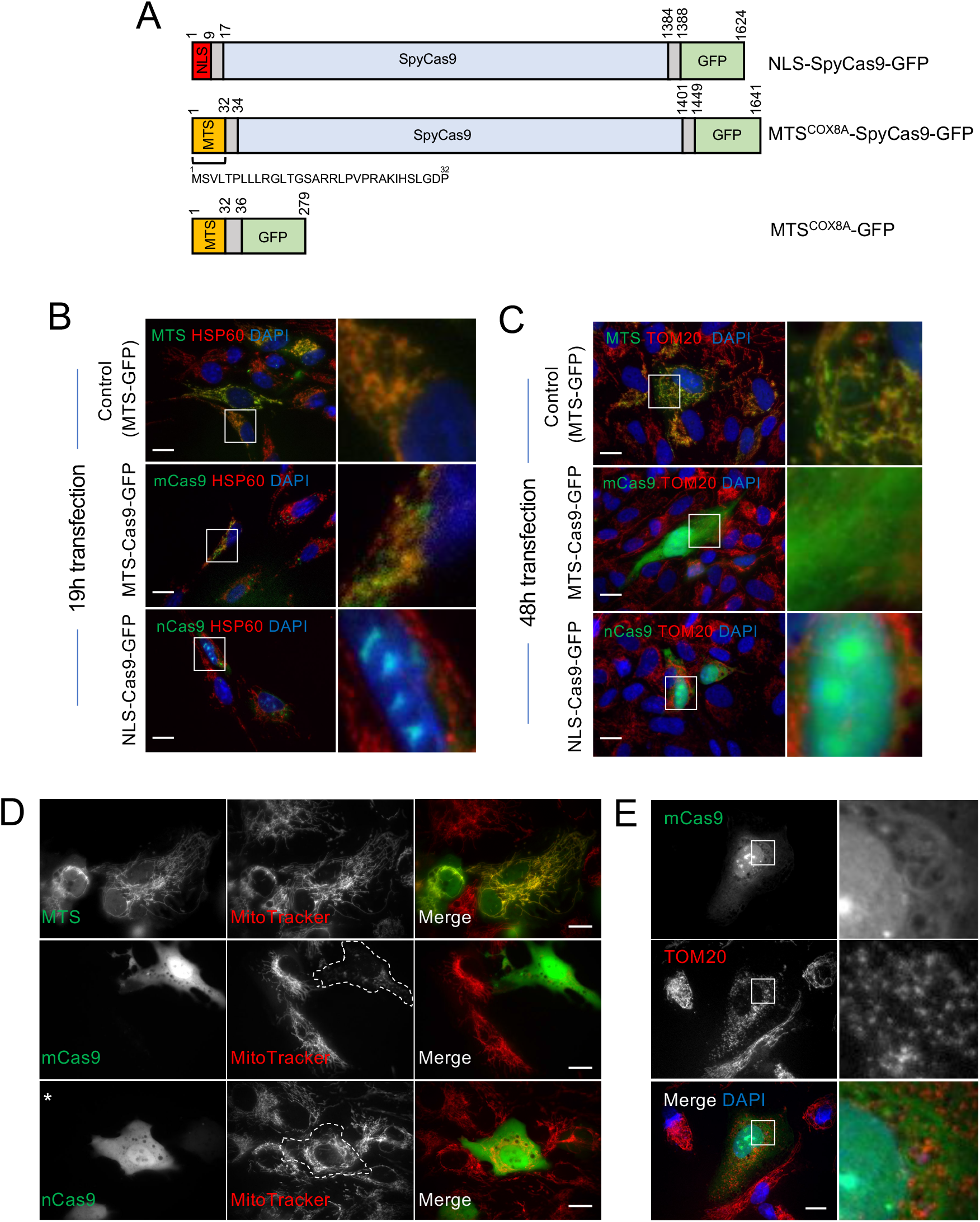
Low SpyCas9 mitochondrial import efficiency is observed using a standard targeting sequence in hTERT-immortalised human RPE1 cells. (A) Diagram representing the constructs used. The nuclear targeting signal is shown in red, the mitochondrial targeting signal in yellow, linkers in grey and GFP in green. (B) In mitoCas9-GFP transfected cells, mitochondria-targeted Cas9 is detected 19 h posttransfection, but the signal is weak. Some cells also have nuclear/cytosolic GFP signal. Scale Bars: 10μm. (C) Cytosolic Cas9 localisation is observed 48h after transfection with mitoCas9-GFP. (D) Loss of mitochondrial membrane potential (absence of Mitotracker fluorescence) and (E) fragmented mitochondria are observed in mitoCas9-GFP-expressing cells. (*) = 10-fold lower exposure time was applied. Scale Bars: 10 μm.

Cells transiently transfected with MTS-GFP showed mitochondrial GFP localisation confirming the ability of the COX8A MTS to localise proteins to the mitochondria under our experimental conditions (**Figure 3B**). Using MTS-SpyCas9-GFP, some GFP signal was observed in mitochondria of RPE1 cells following a 19-hour transfection (**Figure 3B**). However, the signal was weak, and some cells still showed nuclear and cytosolic GFP signal. As expected, NLS-SpyCas9-GFP was mostly localised to the nucleus. Over longer MTS-Cas9-GFP expression periods (up to 48 hours), mitochondrial physiology and morphology were disrupted, and the GFP signal became progressively cytosolic (**Figure 3C**). The impact of MTS-Cas9-GFP expression on mitochondrial health was supported by live imaging of transiently transfected cells stained with Mitotracker Red (**Figure 3D**). At early stages of transfection, and in cells expressing low levels of MTS-Cas9-GFP, mitochondrial networks appeared healthier than in cells expressing high levels of MTS-Cas9-GFP (white dashed line). This is indicated by mitochondria becoming fragmented (**Figures 3D and E**) and by the loss of mitochondrial membrane potential (**Figure 3D**), determined by the absence of MitoTracker fluorescence. In contrast, mitochondria in the NLS-Cas9-GFP expressing cells appeared healthier, despite showing higher expression levels (**Figure 3D**, asterisk indicates lower exposure time). Together these results indicate that SpyCas9 was not being efficiently targeted to mitochondria and, where targeted, is likely disruptive to mitochondrial homeostasis.

### Comparative analysis of the mitochondrial import of various CRISPR-Cas systems

To improve CRISPR nuclease targeting efficiency and avoid mitochondrial damage, we appended alternative MTSs to SaCas9, LbCas12a, and AsCas12a. As with SpyCas9, there was no expectation that any of these proteins would imported into mitochondria in their native forms, but as there might be potential intrinsic features that would affect this process, we first assessed the probability of each being imported into mitochondrial in their native forms. **Figure 4A** shows the domain organisation of SpyCas9, SaCas9 and LbCas12a, highlighting that the overall structure and organisation of Cas9 and Cas12a are quite different [52]. LbCas12a has a RuvC-like endonuclease domain that is similar to the RuvC domain of Cas9; however, LbCas12a does not have an HNH endonuclease domain. One of the main structural differences is that the N-terminus of LbCas12a adopts a mixed α/β structure that is distinct to the N-terminal α-helical recognition lobe of Cas9 (**Figure 4A**). Thus, this part of the protein might influence import to mitochondria. Moreover, using a bioinformatic prediction tool for identifying putative mitochondrial pre-sequences and cleavage sites [53], LbCas12a was suggested to have a much higher predisposition to be imported to mitochondria than the other CRISPR protein variants (**Figure 4B**).

**Figure 4.**
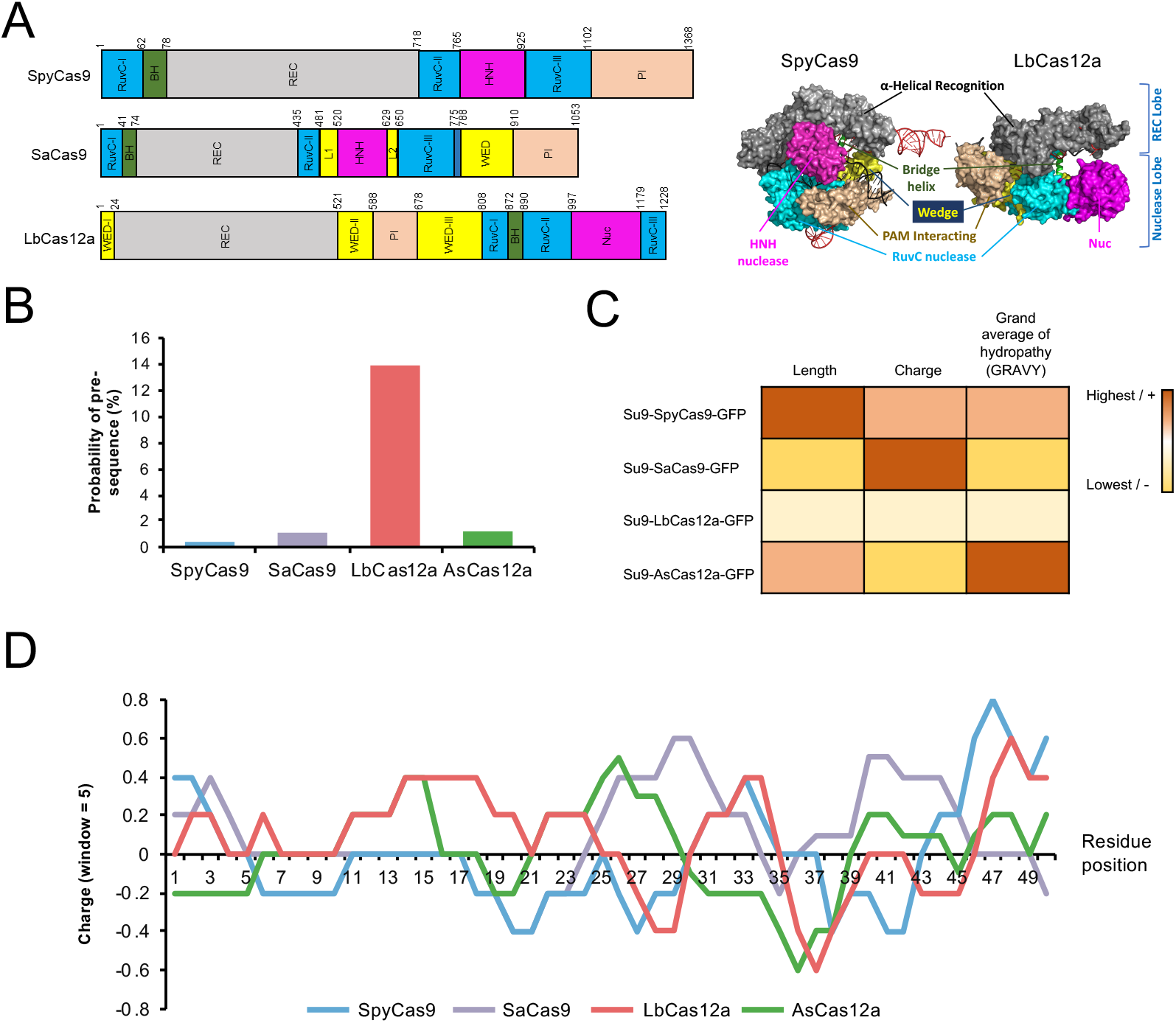
Prediction of the efficiency of the different CRISPR Cas nucleases to be imported into mitochondria. (A) Left-hand panel: schematic diagram of the domain organization of SpyCas9, SaCas9 and LbCas12a. Right-hand panel: overall structures of the SpyCas9-crRNA and LbCas12a-crRNA complexes. Individual protein domains are coloured according to the scheme in the left-hand panel. (B) Probability of containing a N-terminal mitochondrial targeting signal for SpyCas9, SaCas9, LbCas12a and AsCas12a using the MitoFates prediction method [53]. (C) Comparison of parameters that would predict the import efficiency of Su9-SpyCas9/SaCas9/LbCas12a/AsCas12a-GFP (See **Table 3**). (D) Linear charge distribution of the first 50 amino acids of SpyCas9, SaCas9, LbCas12a and AsCas12a calculated using the EMBOSS:charge Bioinformatic tool (charge averaged over 5 residue window) (http://www.bioinformatics.nl/cgi-bin/emboss/charge).

Other parameters that would predict the efficiency of mitochondrial import of a construct were also analysed (**Figure 4C, Table 3**). In terms of the MTS, together with the above-mentioned COX8a example, we compared the 69-residue MTS from *Neurospora crassa* ATPase subunit 9 (Su9) [54, 55], and a cryptic 42-residue MTS from mammalian ATG4D, located immediately downstream of a caspase cleavage site [56]. A more positively charged targeting sequence has been found to enhance import via the mitochondrial membrane potential [57]. Additionally, lengthening an MTS without changing its linear charge density has been shown to increase import [58]. Thus, Su9 would be the preferred choice. When analysing their size, relative charge and hydrophobicity, LbCas12a and SaCas9 were predicted to be imported more efficiently, as they are smaller and less hydrophobic than AsCas12a or SpyCas9. This could explain the poor mitochondria targeting efficiency shown with SpyCas9 (**Figure 3**). Another parameter could be the linear charge distribution of the N-termini (**Figure 4D**), since this will be the first region to be imported. LbCas12a is more positively charged at the N-terminus than SaCas9 (first 24 amino acids), so it could be more efficiently imported, whereas the latter is significantly smaller and has an overall more positive charge.

Mitochondrial import of the three versions of each CRISPR protein with either COX8a, Su9, or ATG4D MTS was tested (**Figure 5A**; showing LbCas12a as an example). All of the tested MTSs showed mitochondrial localisation by fluorescence microscopy as GFP fusion control constructs. Using the compact SaCas9, targeting efficiency was clearly improved and mitochondria looked healthier, but relatively few cells showed mitochondria targeted Cas9 and only a small proportion of the protein was localised to mitochondria (in most transfected cells, GFP was detected in the nucleus likely corresponding to mis-targeting Cas9 or free GFP) (**Figure 5B**). However, in the case of LbCas12a, using any of the MTSs, the protein was reliably localised to mitochondria and the organelles looked healthy by fluorescence microscopy. In cells expressing COX8a-LbCas12a/AsCas12a-GFP or Su9-AsCas12a-GFP, some cytosolic signal was also observed, possibly due to protein mis-targeting. In contrast, using either Su9-LbCas12a or ATG4D-LbCas12a/AsCas12a, almost all transfected cells showed mitochondrial targeting and healthy mitochondria (**Figure 5B**). This mitochondrial colocalisation was confirmed by a semiquantitative analysis of the images using fluorescence profiles of GFP and MitoTracker Red signals taken along a 150px line (**Figure 5B**). A specific antibody against LbCas12a was also used to confirm LbCas12a mitochondrial localisation in transiently transfected RPE1 cells (**Figure 5C**). Optimal expression was observed in lysates of RPE1 cells transfected with Su9LbCas12a-GFP (24h post-transfection) (**Figure 5D**). A band was also detected in the COX8a-GFP sample fractionated by 8% SDS-PAGE, with a size that does not correspond to any of the constructs tested. Two different constructs were made as control versions of the mitochondria-targeted LbCas12a: Su9LbCas12a without the GFP tag and LbCas12a-GFP without the MTS. As expected, Su9LbCas12a was localised to mitochondria in either RPE1 cells or WT/MELAS cybrids and the non-targeted version of LbCas12a was found to be cytosolic in RPE1 cells (**Figures S2A and B**).

**Figure 5.**
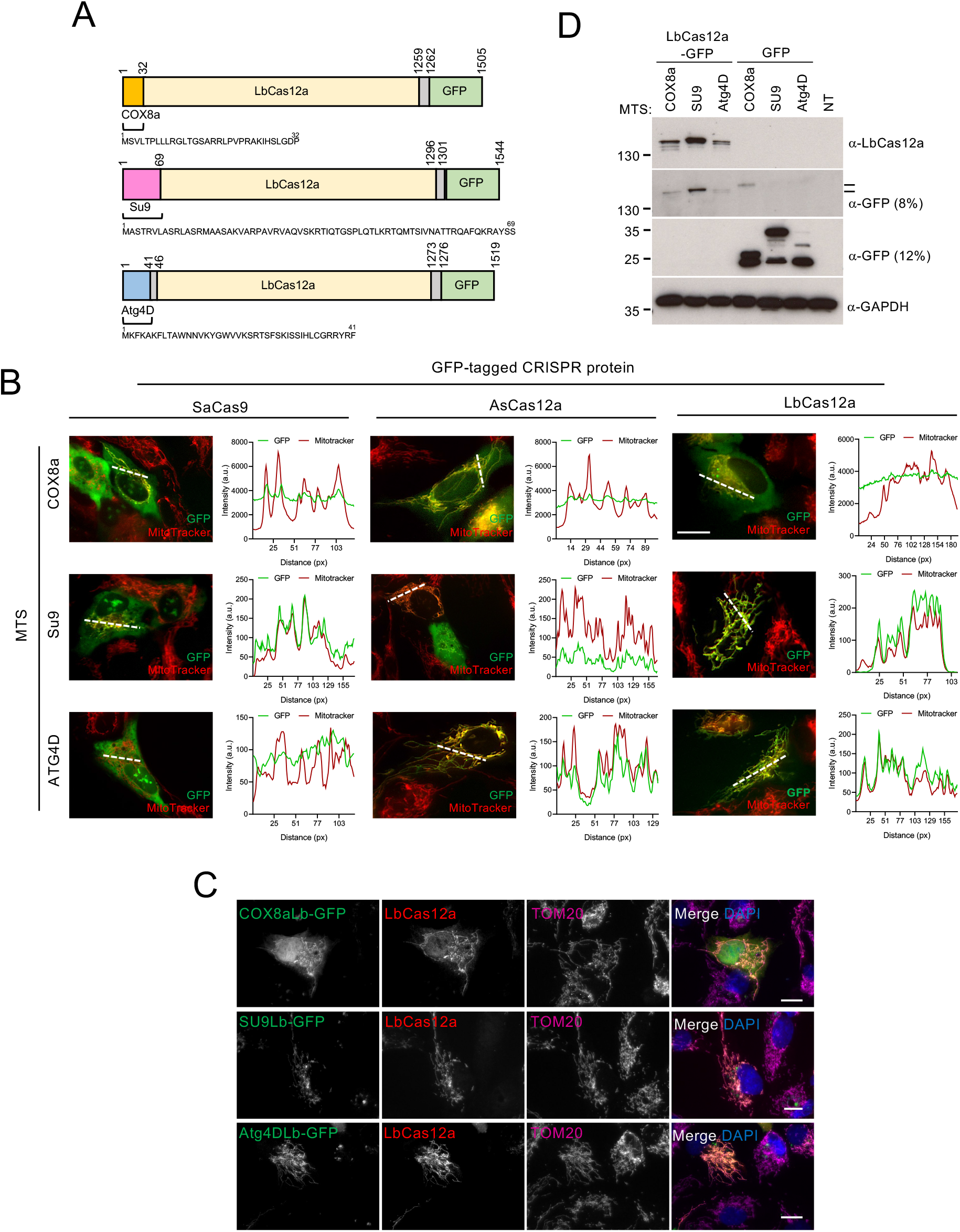
Su9LbCas12a-GFP showed the highest mitochondrial targeting efficiency and optimal expression levels in hTERT-immortalised human RPE1 cells. (A) Diagram representing the constructs used. The mitochondrial targeting sequence COX8a is shown in yellow, Su9 in pink, ATG4D in blue, and linkers in grey. (B) Expression and localisation of LbCas12a-GFP, AsCas12a-GFP, and SaCas9-GFP with various N-terminal mitochondrial targeting sequences (MTSs) in hTERT-immortalised human RPE1 cells. Scale bars: 10μm (see **Figure 5B** for semi-quantitative companion data). (C) Mitochondria-targeted LbCas12a is detected in mitoLbCas12a-GFP transfected cells using any of the MTSs tested. Scale bars: 10μm. (D) Optimal expression is observed in lysates of RPE1 cells transfected with Su9LbCas12a-GFP following 24h transfection.

Overall, our fluorescence data correlate with the bioinformatic predictions for the mitochondrial import of these CRISPR variants. We can therefore conclude that LbCas12a traffics most efficiently to mitochondria, in particular using the Su9 MTS.

### Assessment of LbCas12a-based heteroplasmy purification in MELAS cybrid cells: considerations for MitoCRISPR tool development

The MitoCRISPR attempts to date have measured changes in mtDNA copy number [25, 27, 29]; however, a better way to validate targeted nuclease activity is to demonstrate heteroplasmy purification, as seen, for example, with MitoTALENs [13]. In this study, we used cybrid cell-line versions of a mitochondrial encephalomyopathy, lactic acidosis and stroke-like episodes (MELAS) syndrome mutation as a possible model system to test the development of Cas12a-based MitoCRISPR tools. MELAS is one of the most common mitochondrial disorders. It is associated with neurological symptoms and other secondary manifestations such as depression, cardiomyopathy and diabetes mellitus [5][6][7]. Although this syndrome can be caused by different mutations, 80% of patients harbour a transition of adenine to guanine at the 3243 position in the MT-TL1 gene (tRNALeu (UUR)) of mtDNA. The mutation leads to defects in mitochondrial protein synthesis and respiratory chain function [8, 9]. This MELAS mutation has the added practical advantage in this study of allowing rapid identification of WT and mutant mtDNA levels by PCR-RFLP, as the mutation creates a novel restriction site for the enzyme, ApaI.

During our attempts to establish a working MitoCRISPR system using Su9LbCas12a-GFP with nontargeted and 5′RP gRNAs (based on previous reports of MitoCRISPR-type activity using non-targeted gRNAs [27, 29]), we encountered several practical difficulties that highlight the challenges of developing MitoCRISPR as a tool (detailed in **Figure 6, Figure S7,S8**): (1) despite few obvious morphological changes, mitochondria in Su9LbCas12a-GFP stable cell-lines have reduced respiratory capacity; (2) as proteins are imported into mitochondria in their unfolded state, this precludes the use of assembled ribonuclear particles; (3) the limited availability of suitable PAM sequences (a limitation of any CRISPR system, but particularly problematic for heteroplasmy purification that must be targeted to specific mutations); (4) the lack of specificity towards mutant vs. wild-type mtDNA, due to inherent insensitivity of LbCas12a to mismatches; (5) inherent crRNA processing by LbCas12a, a feature that could prove useful for limiting the persistency of *in situ* CRISPR activity; (6) poor reproducibility in functional MitoCRISPR assays. In the following sections, we present our findings with respect to these limitations, along with attempts to demonstrate LbCas12a-mediated MitoCRISPR activity in cybrid cells.

**Figure 6.**
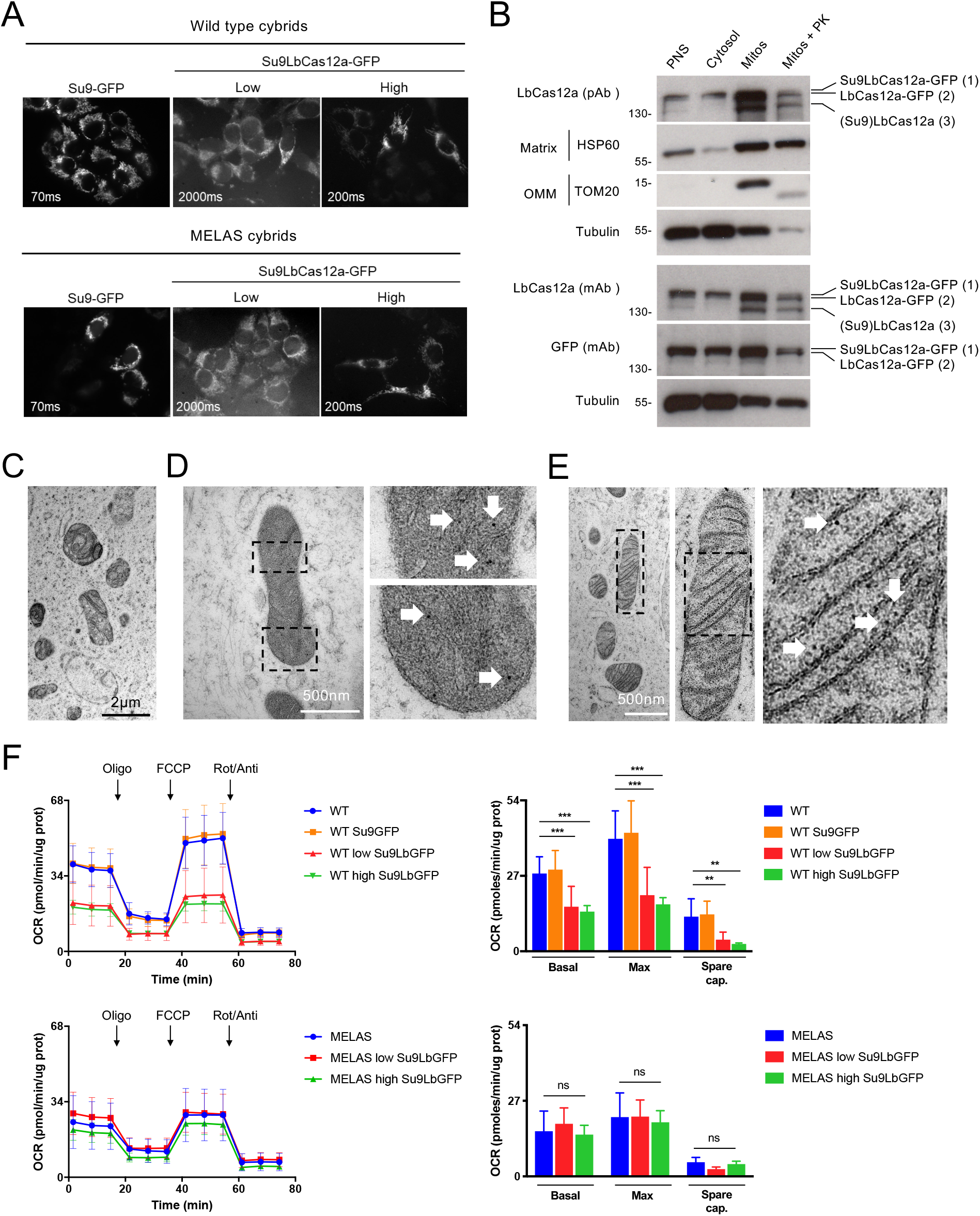
LbCas12a is imported through TOM/TIM route and is localised to the mitochondrial matrix in WT or MELAS cybrids expressing Su9LbCas12a-GFP. (A) WT and MELAS cybrids stably expressing Su9-GFP or Su9LbCas12a-GFP with low or high expression levels. (B) Subcellular fractionation of WT cybrids transiently expressing Su9LbCas12a-GFP. Top panel: band pattern using polyclonal anti-LbCas12a. Bottom panel: band pattern using monoclonal anti-LbCas12a and anti-GFP. (C-D) ImmunoEM images of mitochondria in WT cybrids stably expressing Su9LbCas12a-GFP stained with anti-GFP (C, D) or anti-LbCas12a (E) and 10nm gold secondary antibodies. Arrows indicate gold particles. (F) Mitochondrial respiration profile of WT and MELAS cybrids stably expressing Su9LbCas12a-GFP measured using a XF Cell Mito Stress Test (Seahorse). Student’s t test: **p<0.01; ***p<0.001 vs control.

#### Mitochondrial targeting of Su9LbCas12a-GFP in cybrid cell-lines for MitoCRISPR

Based on ApaI-mediated digestion, the heteroplasmy load of the cybrids used in this project was 100% for both wild-type (WT) and MELAS (**Figure S3A**). Su9LbCas12a-GFP and Su9-GFP were stably expressed in the cybrids by lentiviral infection and FACs sorting (**Figure 6A, Figure S3B-D**). Mitochondrial-targeted LbCas12a was detected at different levels in WT or MELAS cybrids stably transfected with Su9LbCas12a-GFP. LbCas12a localisation to the mitochondrial matrix was confirmed by cell fractionation (**Figure 6B**), TOM/TIM import blockers (**Figure S4**), and immunoEM (**Figure 6C-E**). In the cell fractionation experiments, mitochondria were isolated from WT cybrids transfected with Su9LbCas12a-GFP and treated with proteinase K to test whether mitochondria-targeted LbCas12a was protected from degradation (and therefore had been fully imported). Following the proteinase K treatment, both the mitochondrial matrix marker HSP60 and LbCas12a, but not the OMM marker TOM20, were retained (**Figure 6B**). Moreover, three LbCas12a bands were observed in the mitochondrial fraction using either polyclonal or monoclonal antibodies. These comprised: (1) the precursor form; (2) the mature protein; and (3) a product likely corresponding to Su9LbCas12a separated from GFP (as this band was not detected with the antibody recognising GFP). Notably, the precursor:mature LbCas12a ratio appeared to change following proteinase K treatment, such that only the precursor form was accessible and susceptible to degradation. WT cybrids stably expressing Su9LbCas12a-GFP were processed for immunoEM using anti-GFP (**Figures 6C and D**) or anti-LbCas12a (**Figure 6E**) primary antibodies, and 10 nm gold secondary antibodies. Mitochondria with damaged or missing cristae were observed in many of these cells (**Figure 6C**); however, focusing on mitochondria of healthy appearance, Su9LbCas12a-GFP was found localised to the mitochondrial matrix (**Figure 6D, E**). Import of LbCas12a to the mitochondrial matrix through TOM/TIM import route was confirmed by the mitochondrial import blockers (MitoBloCK: MB-10 (TIM23 inhibitor); MB-12 (DECA: TIM23 and ATP synthase inhibitor)) (**Figure S4**). As expected, mis-localisation of the Su9LbCas12a fusion protein from mitochondria to the cytosol was observed in RPE1 cells (**Figure S4A**) or WT/MELAS cybrids (**Figure S4B, C**) treated with 20μM MB-10 or 10μM MB-12 for 24 hours.

#### The impact of LbCas12a mitochondrial import on mitochondrial respiration

The bioenergetic profile of cybrids stably expressing Su9LbCas12a-GFP was measured using the Seahorse XF Cell Mito Stress Test (**Figure 6F**). As expected, WT cybrids, but not MELAS cybrids, showed normal mitochondrial respiration (oxygen consumption rate, OCR) profiles. However, OCR measured in WT cybrids stably expressing mitochondrial-localised LbCas12a at either low or high levels was significantly lower (**Figure 6F, top panel**). This was not observed with the control stable line expressing Su9-GFP. The stable MELAS cybrids did not show a normal mitochondrial respiration profile in any tested condition. These data indicate that prolonged expression of Su9LbCas12a-GFP, whilst not adversely and grossly affecting mitochondrial morphology, does impact on mitochondrial functionality at the level of mitochondrial respiration. We can therefore conclude that LbCas12a is localised to the matrix as confirmed by cell fractionation or EM, but that mitochondrial respiration is affected in cybrids stably expressing Su9Cas12a-GFP.

#### Requirement for unfolding of LbCasl2a during mitochondrial import

Any effective MitoCRISPR system would need to consider the mode of delivery of both CRISPR protein and gRNA. In this study, we have focussed on expressing CRISPR proteins on plasmids and viral vectors. We also considered whether native CRISPR proteins appended with MTSs could be imported into mitochondria as pre-assembled ribonuclear particles for MitoCRISPR. **Figure S5** shows that import into isolated yeast mitochondria of recombinant Su9-LbCas12a-myc requires chemical denaturation, as no product is detected in proteinase K-treated samples (with or without intact proton motive force). To test for possible import of mitochondrial-targeted Casl2a post-translation in living cells, we treated WT cybrids stably expressing Su9-LbCas12a-GFP with MB-10 for 24 hours (to block import), then washed out the drug and incubated cells for up to a further 16 hours in the presence of cycloheximide (to block new protein expression). Although the bulk of the GFP symbol remained cytosolic, a minor fraction of the protein produced before translation inhibition and after the release of the import block co-localised with TOM20-labelled mitochondria. This might be indicative of partial import of pre-folded Cas12a (**Figure S6**).

#### Analysis of mitochondrial LbCas12a cleavage activity and mtDNA levels in MELAS cybrids

The next step was to design Cas12a crRNAs specific for the MELAS mutation. Two different crRNAs were used to target the *MT-TL1* gene comprising the MELAS mutation (**Figure S7A**). Each of them targeted a different DNA strand, and both were tested as untargeted or mitochondrial-targeted versions. First, the DNA cleavage activity of LbCas12a using the four different gRNAs was assessed by cleavage assays *in vitro* (**Figure S7B**). In these experiments, WT or mutant MT-TL1 DNA was cloned into a plasmid (pSP1) which was cut with a similar cleavage efficiency using any of the designed crRNAs. However, the cleavage appeared to be nonspecific as LbCas12a targeted both WT and mutant DNA containing the MELAS mutation (ApaI was used to distinguish the WT and mutant target sequences). The unspecific binding could be due to the insensitivity of Cas12a to mismatches at certain locations in the spacer ([59], https://www.biorxiv.org/content/10.1101/696393v1), or due to a recently reported lack of specificity of Cas12a under some conditions (https://www.biorxiv.org/content/10.1101/657791v1). It should be noted that the cleavage activity is likely to be independent of a 5′ modification since the 5′RP crRNA was found to be processed by LbCas12a *in vitro* (**Figure S7C**). This feature of Cas12a is potentially useful since a 5′ modification could be made to a crRNA to allow efficient import, and upon RNP assembly in the matrix, the crRNA would be processed so decreasing the likelihood of possible off-target effects.

#### Testing Su9LbCas12a-GFP for evidence of MitoCRISPR activity in MELAS cybrids

Despite its apparent lack of specificity, we decided to test the functionality of the Cas12a-crRNA system inside mitochondria. We compared three versions of LbCas12a: WT LbCas12a; a PAM interacting (PI) domain mutant (lacking the PI domain); or a RuvC inactive version containing the point mutation D832A (dLbCas12a) (**Figure S8A**). We opted to use transient transfection of Cas12a proteins and crRNAs to avoid the functional decline observed in mitochondria over time in stable cell-lines. Both WT and mutant forms of Su9-LbCas12a-GFP localised strongly to mitochondria in MELAS cybrids co-transfected with a control (PLKO.1) plasmid (**Figure S8B**). Next, mtDNA levels were analysed in MELAS cybrids transiently co-transfected with unmodified crRNAs (crRNA(1)/(2)) or 5′RP crRNAs and mitochondria-targeted Su9-LbCas12a-GFP (**Figure S8C-H**). crRNA transcription was achieved by cloning the corresponding crRNA sequence into a PLKO.1 vector containing a U6 promoter.

Transcriptional activity of the transfected PLKO.1 plasmids was evaluated from crRNA levels using qRT-PCR in MELAS cybrids co-transfected with WT LbCas12a (Su9LbCas12a-GFP) (**Figure S8C**). Since we used homoplasmic MELAS cybrids as the model system, a reduction in mtDNA copy number was expected following CRISPR nuclease activity inside mitochondria. In fact, mtDNA double-strand breaks have been demonstrated to not be repaired, but rather the broken mtDNA is eliminated, probably by mitochondrial nucleases [5]. To evaluate this, COXII DNA levels were measured by q-PCR following 48 hour cotransfection of cybrids with WT or mutant Su9LbCas12a-GFP and the corresponding crRNA or empty PLKO.1 control vector (**Figure S8D**). An unexpected increase in mtDNA levels was observed after transfection of MELAS cybrids with WT Su9LbCas12a-GFP and crRNA(2) or 5′RP crRNA(2). This effect was not observed using any of the mutant forms of Su9LbCas12a-GFP or using the gRNA(1)/5′RP(1) RNA versions (**Figure S8D**), suggesting the nuclease activity might be necessary to observe this effect. Similar crRNA transcription levels were observed using any of the protein variants after co-transfection of MELAS cybrids with 5′RP cRNA(2) and WT or mutant forms of Su9LbCas12a-GFP (**Figure S8E**). This indicates that the lack of effect on mtDNA levels using mutant forms of Su9LbCas12a-GFP was not due to a lower crRNA transfection or expression efficiency. Moreover, MT-TL1 and COXII mRNA levels also increased after co-transfection with WT Su9LbCas12a-GFP and crRNA(2) (**Figures S8F,G**). In a further validation experiment to confirm the increase in COXII DNA levels after co-transfection with WT Su9LbCas12a-GFP and gRNA(2), no significant changes in mtDNA levels were observed (**Figure S8H**). This indicates that the mitochondria targeted CRISPR/Cas12a system effects are, at best, variable.

## DISCUSSION

Given the impact that technology for the efficient manipulation of mtDNA would have on the study of mtDNA and disease, it is not surprising that there have been several published efforts to develop MitoCRISPR techniques. However, none have yet described a tool that is as efficient as the best ZFN and TALEN examples. The most successful iteration to date has reported a depletion of mtDNA copy number [25], but CRISPR-mediated heteroplasmy purification, the gold-standard for mtDNA editing, has yet to be achieved. This work here aimed to provide a careful validation of MitoCRISPR toolkit parts.

Despite being used in all the previous studies, mitochondria targeted SpyCas9 (COX8A-Cas9) showed poor mitochondrial localisation in transiently transfected mammalian cells as analysed by immunofluorescence. Moreover, cells expressing mitochondria targeted SpyCas9 suffered a dramatic deterioration in mitochondrial morphology and function, which is likely to impact on cell viability. Overexpression and mitochondrial import of this large protein is likely to be disruptive to mitochondrial morphology/physiology. In contrast, LbCas12a was successfully expressed and localised to the mitochondrial matrix using several MTSs as measured by immunofluorescence, western blotting and immunoEM, with the most successful MTS being Su9. However, careful analysis of mitochondrial LbCas12a import and cellular impact using cell-lines stably expressing Su9-Cas12a-GFP showed evidence for mitochondrial damage caused by the prolonged expression of mitochondria-targeted LbCas12a. Therefore, if the mitochondria targeted CRISPR/LbCas12a system were to be used further, transient expression would likely be needed to prevent mitochondrial disturbance. However, a more significant problem in using Cas12a is possible off-target cleavage activity. In contrast to the very reliable nuclease gene knockouts of classical CRISPR, heteroplasmy purification requires that the targeting occurs exactly at the mutated sequence. With the limitations of PAM availability, protospacer sequence choice could be limited to a single sequence. Consequently, if mismatches are tolerated by Cas12a, it will be difficult to achieve specificity.

The obvious difference in mitochondrial import efficiency observed for SpyCas9 and LbCas12a could be explained by various characteristics of the proteins overviewed in this work. The overall domain organisation of these enzymes is significantly different, particularly with respect to their N-terminal secondary structures which are typically key for successful protein import into mitochondria. Furthermore, amino acid bias or differences in overall peptide charge of imported proteins also needs be considered, as favourable interactions with the TOM/TIM machinery will encourage mitochondrial import. This is consistent with LbCas12a being smaller and less hydrophobic than SpyCas9, as well as having a more positively charged N-terminus (first 24 amino acids), which would also explain the higher mitochondria targeting efficiency shown for Cas12a. Moreover, MTS accessibility is important for protein import, and it may be that the peptide is sub-optimally presented on Cas9. The use of alternative MTSs or introducing a flexible linker between the MTS and Cas9 structure in conjunction with directed *in silico* modelling, might ameliorate this. An appreciation of those features that favour Cas12a import over Cas9 could also be used to modify Cas9 structure, optimising it for mitochondrial import (for example, using a split Cas9 – [60]).

Alternatively, general rules governing mitochondrial import could be applied when screening the large number of CRISPR-Cas variants in the pangenome [61]. Alternative CRISPR-Cas enzymes may prove more efficient and less disruptive to organelle health.

Our data on RNA import add to other evidence for mitochondrial localisation of gRNAs without the requirement for targeting modifications [7, 25, 27]. It should be noted that our assays do not explicitly demonstrate import, but rather RNAse protection and/or co-fractionation. For the protein import assays, evidence of delivery to the matrix is ascertained by a change in the protein size as the MTS is proteolytically removed (as seen in our *in vitro* Cas12a import assay; **Figure S5**). However, there are not any reports of an equivalent RNA aptamer processing activity in mitochondria to use as a functional readout. Further careful examination of mitochondrial compartments would be needed to prove whether the gRNA is matrix-located. A limitation of the RP-facilitated import of Cas9 gRNAs for mtDNA editing is that once bound to Cas9, 5’RP modified gRNAs only support formation of small R-loops and completely block DNA cleavage (Mullally and Szczelkun, unpublished observations). The processing of 5’ modifications by Cas12a removes any potential inhibition and is thus an additional advantage of the Type V system over Cas9.

Data presented here indicate that in the presence of an active mitochondria targeted Cas12a and crRNA, it is possible to observe an increase in the copy number of mtDNA. Given that defects in mitochondria such as dysfunctional mitochondrial dynamics or fragmentation were not observed, the change in mtDNA copy number recorded might be due to a feedback mechanism triggered to (over)compensate for a perceived reduction in mtDNA. However, the lack of reproducibility in this assay currently remains the greatest hurdle in the application of MitoCRISPR. We would argue that any future studies need to use heteroplasmy purification in the absence of significant changes in mitochondrial health as a benchmark for success, and that independent validation should be an important goal.

## ACKNOWLEDGEMENTS

We thank Maggie Hicks, Jia Qi Cheng Zhang and Ciaran Guy for help with preliminary experiments. Su9-EGFP was a gift from David Chan; pLKO.1-TRC was a gift from David Root; pX600-AAV-CMV::NLS-SaCas9-NLS-3xHA-bGHpA and pY016 (pcDNA3.1-hLbCpf1), pY010 (pcDNA3.1-hAsCpf1) were gifts from Feng Zhang; and, MMW1578: CAG-human dLbCpf1(D832A)-NLS-3xHA was a gift from Keith Joung.

