## Supplemental tables and figures for "Variability in mitochondrial import, mitochondrial health and mtDNA copy number using Type II and Type V CRISPR effectors"

### SUPPLEMENTARY FIGURES

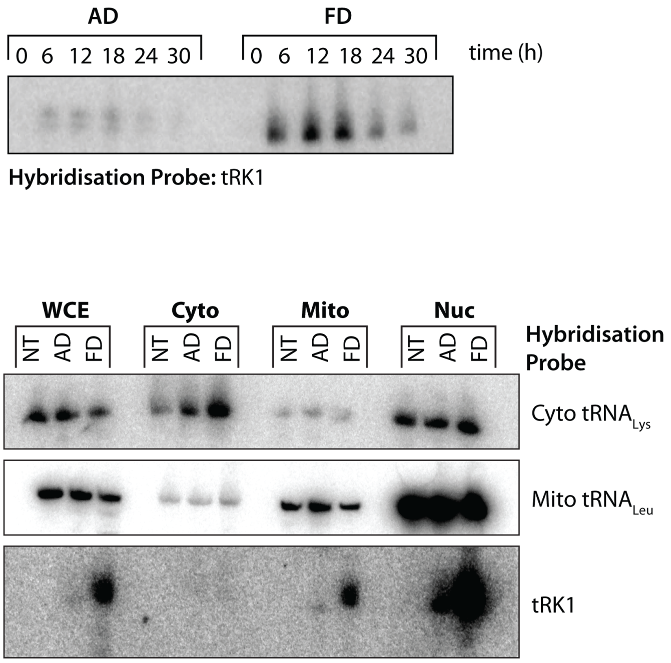


**Figure S1**. (Upper panel) Transfection time course for AD and FD RNA. (Bottom panel) Northern blotting HeLa cells for AD and FD separated by urea-PAGE, transferred to nitrocellulose membrane and probed against tRK1 probe. WCE = whole cell extract; Cyto = Cytoplasm; Mito = Mitochondria; Nuc = Nuclear.

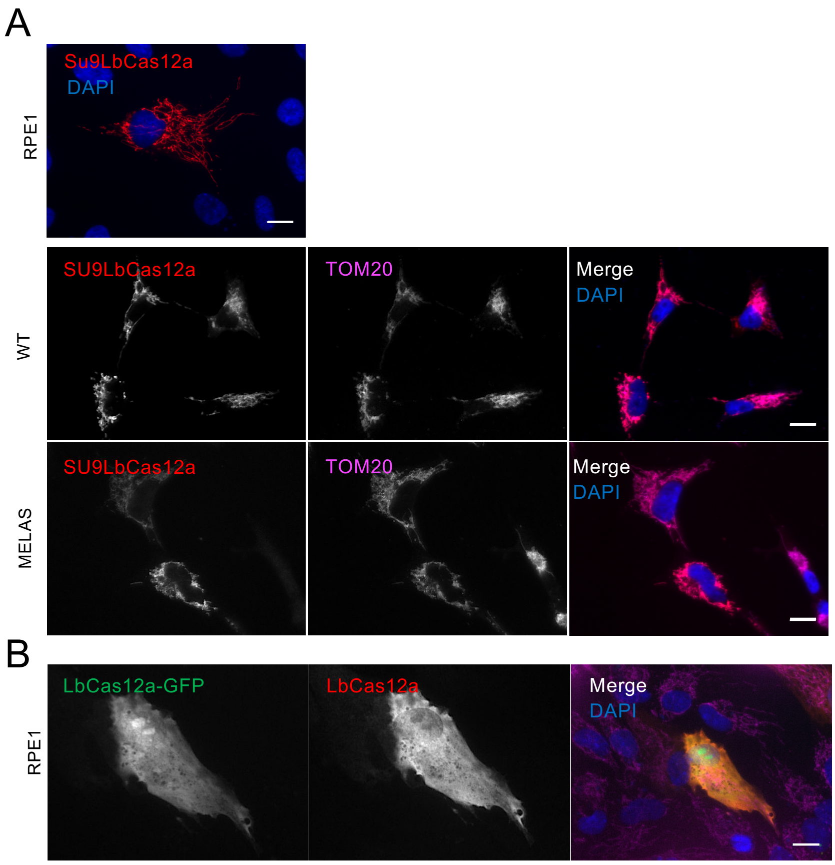


**Figure S2**. Representative images showing (A) Su9LbCas12a localisation to mitochondria in RPE1 cells or WT/MELAS cybrids and (B) untargeted LbCas12a-GFP cytosolic expression.

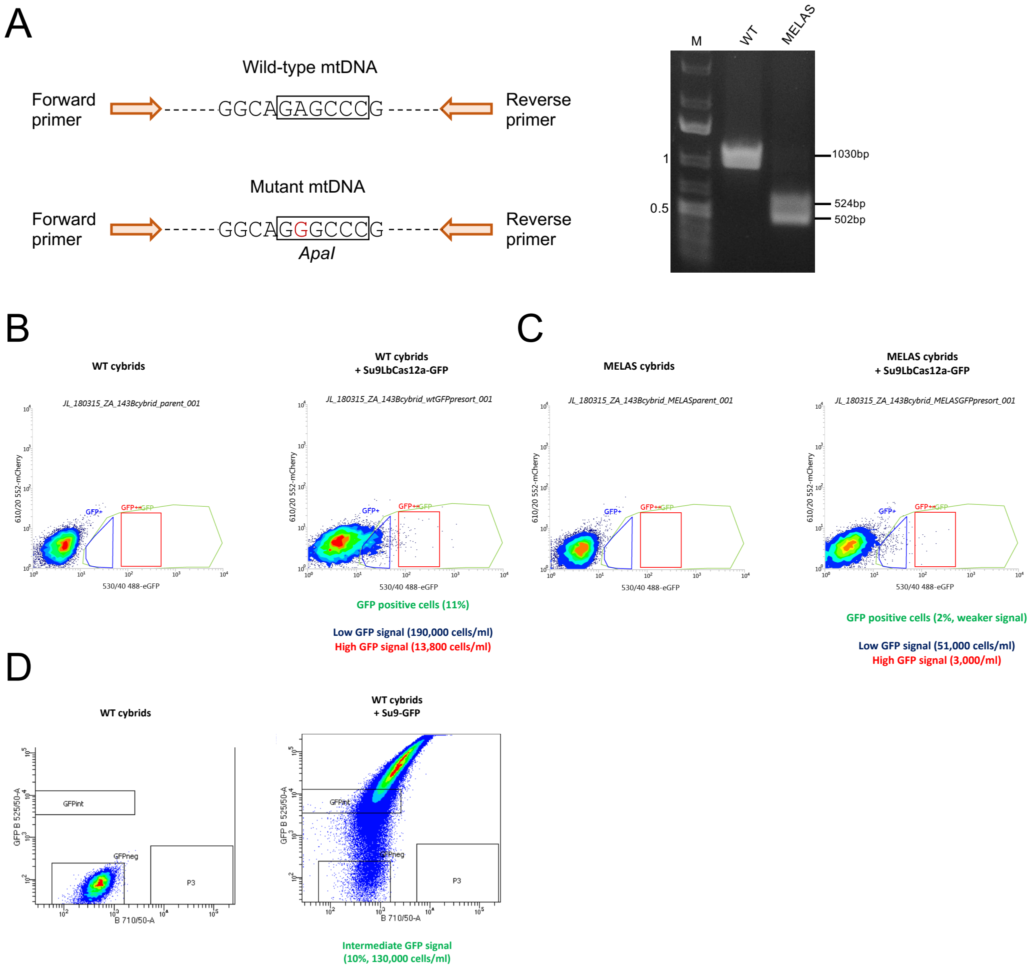


**Figure S3**. (A) Schematic showing the A3243G mutation in MT-TL1 gene containing the ApaI restriction endonuclease site. This mutation was detected by polymerase chain reaction (PCR) amplification of a region of mtDNA containing nt 3243, followed by ApaI digestion and electrophoretic analysis of the resulting fragments. (B-D) Fluorescence activated cell sorting (FACS) of WT and MELAS cybrids expressing Su9LbCas12a-GFP or Su9-GFP.

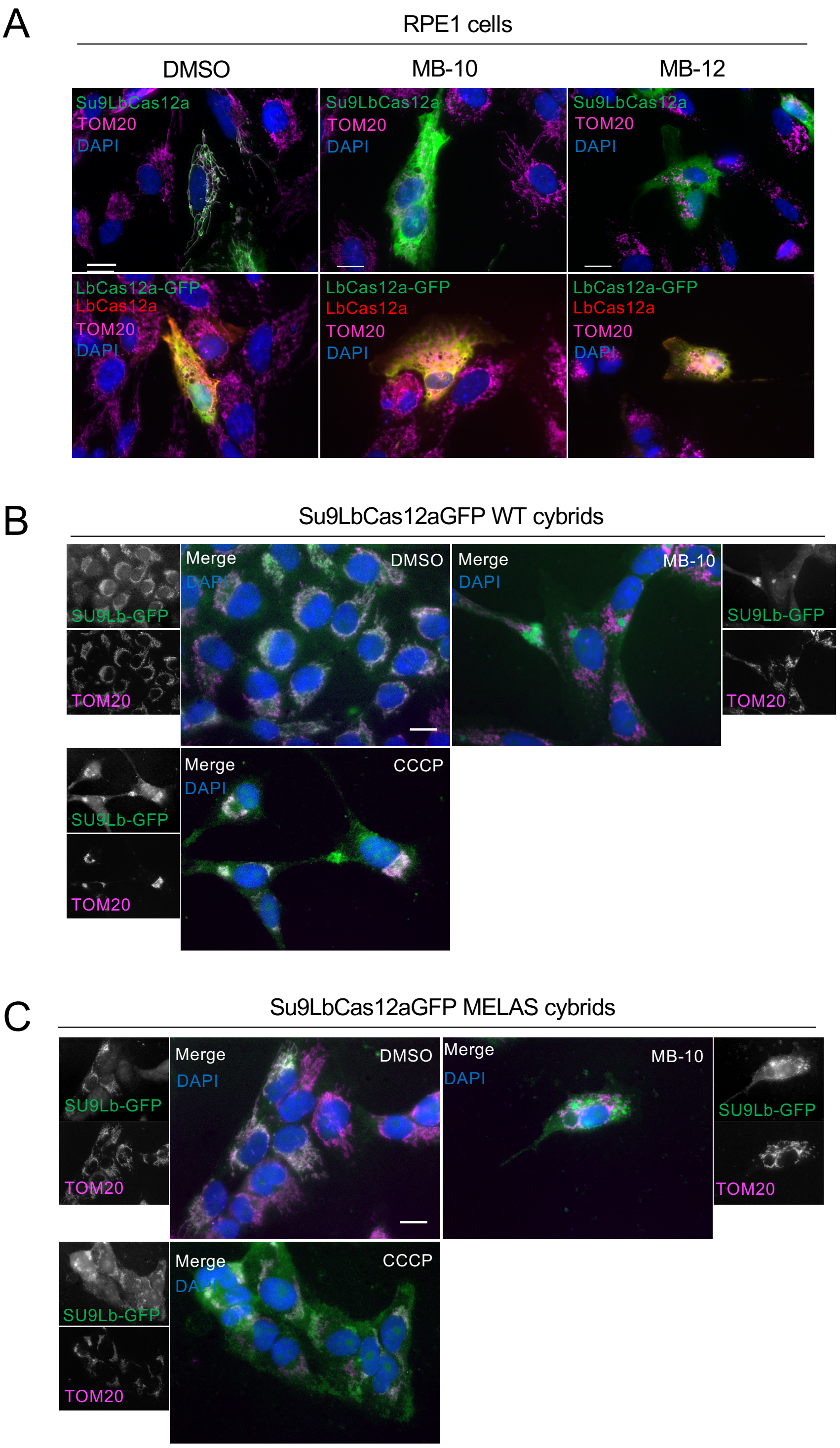


**Figure S4**. (A) Mislocalisation of the Su9LbCas12a fusion protein from mitochondria to the cytosol is observed in RPE1 cells transfected with Su9LbCas12a-GFP following treatment with 20 µM MB-10 or 10 µM MB-12 for 24 h. Scale Bar: 10 µm. (B-C) Mislocalisation of LbCas12a from mitochondria to the cytosol is observed in WT or MELAS cybrids stably expressing Su9LbCas12a-GFP treated with 20 µM MB-10 or 10 µM CCCP for 24 h. Scale Bars: 10 µm.

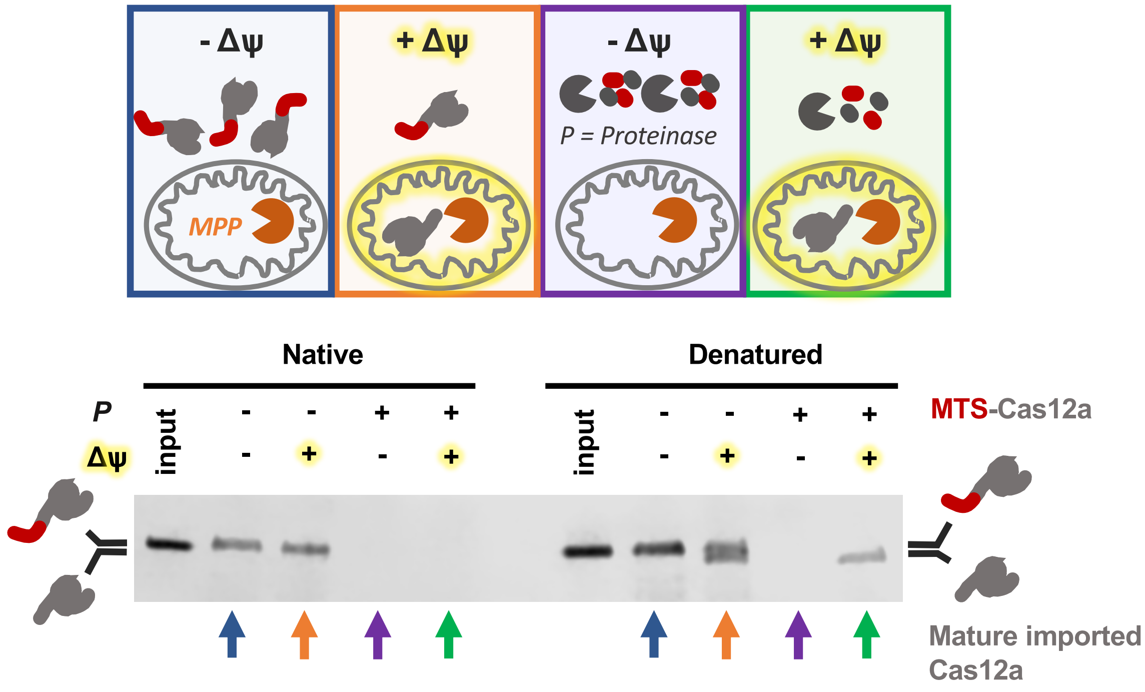


**Figure S5**. Import into isolated yeast mitochondria of purified Su9-LbCas12a requires chemical denaturation. (Upper panel) Cartoon of the assay. Addition of VOA removes the membrane potential needed for import (-Δψ). Imported protein has the MTS removed by the Mitochondrial Processing Peptidase (MPP red Pacman). Treatment of the cells with protease (black Pacman) cleavaes any non-imported protein. (Lower panel) Western blot against the Myc-tag on LbCas12a and the conditions as indicated in the cartoon. In the presence of membrane potential, the urea-denatured protein has two bands, the upper of which is removed by the addition of external protease, indicating import and MTS processing. The native folded protein only gives a single band, all of which is cleaved by the external protease, indicating that import and MTS processing is not occurring.

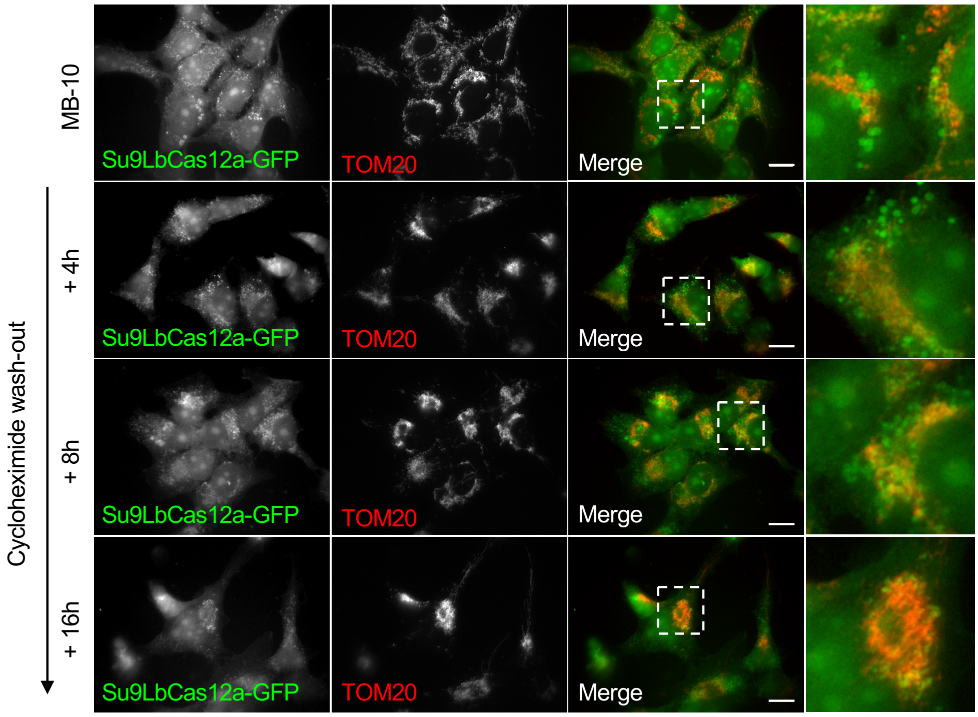


**Figure S6**. Partial mitochondrial reimport of Su9LbCas12a-GFP is observed in stable WT cybrids following incubation with 20 µM MB-10 (24 h) and removal for up to 16 h in the presence of cycloheximide (50 µg/ml). Scale Bars: 10 µm.

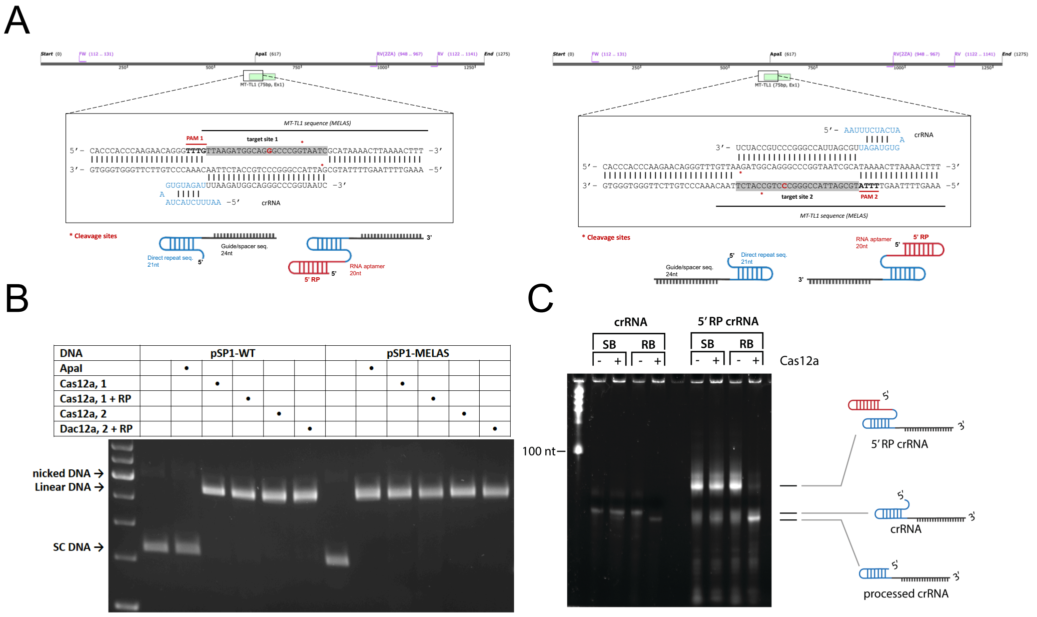


**Figure S7**. Cleavage activity and processing of gRNAs by LbCas12a: Cas12a removes any aptamer from the 5' end. (A) A schematic showing the gRNAs used for functional validation of nuclease activity in the MT-TL1 gene. Two target sequences were tested, and the 5′RP aptamer was used as a mitochondrial targeting signal. (B) Example agarose gel from pSP1 cleavage assay. Unmodified gRNA or 5′RP gRNA was used targeting site 1 or 2 in the MT-TL1 gene. ApaI digestion was used as a control for WT and MELAS DNA. (C) Processing of gRNAs by LbCas12a. 2 μM gRNA or 5′RP gRNA was incubated with 10 μM LbCas12a in 1XRB or 1XSB at 37°C for 1 h. Samples were then separated by urea-PAGE and stained with SYBR-Gold. The data show that crRNAs both with and without the 5′RP aptamer are processed in a magnesium-dependent manner. Unmodified crRNAs undergo a small shift in size, perhaps due to the removal of only a few of nucleotides. As with the SpyCas9 gRNAs, LbCas12a crRNAs also have an additional 5′ guanine resulting from *in vitro* transcription, so it is likely that this extra nucleotide is what is being processed here. 5′RP crRNAs incubated with Cas12a in 1XRB are processed to the same size as unmodified crRNAs, presumably the result of the RP hairpin being removed.

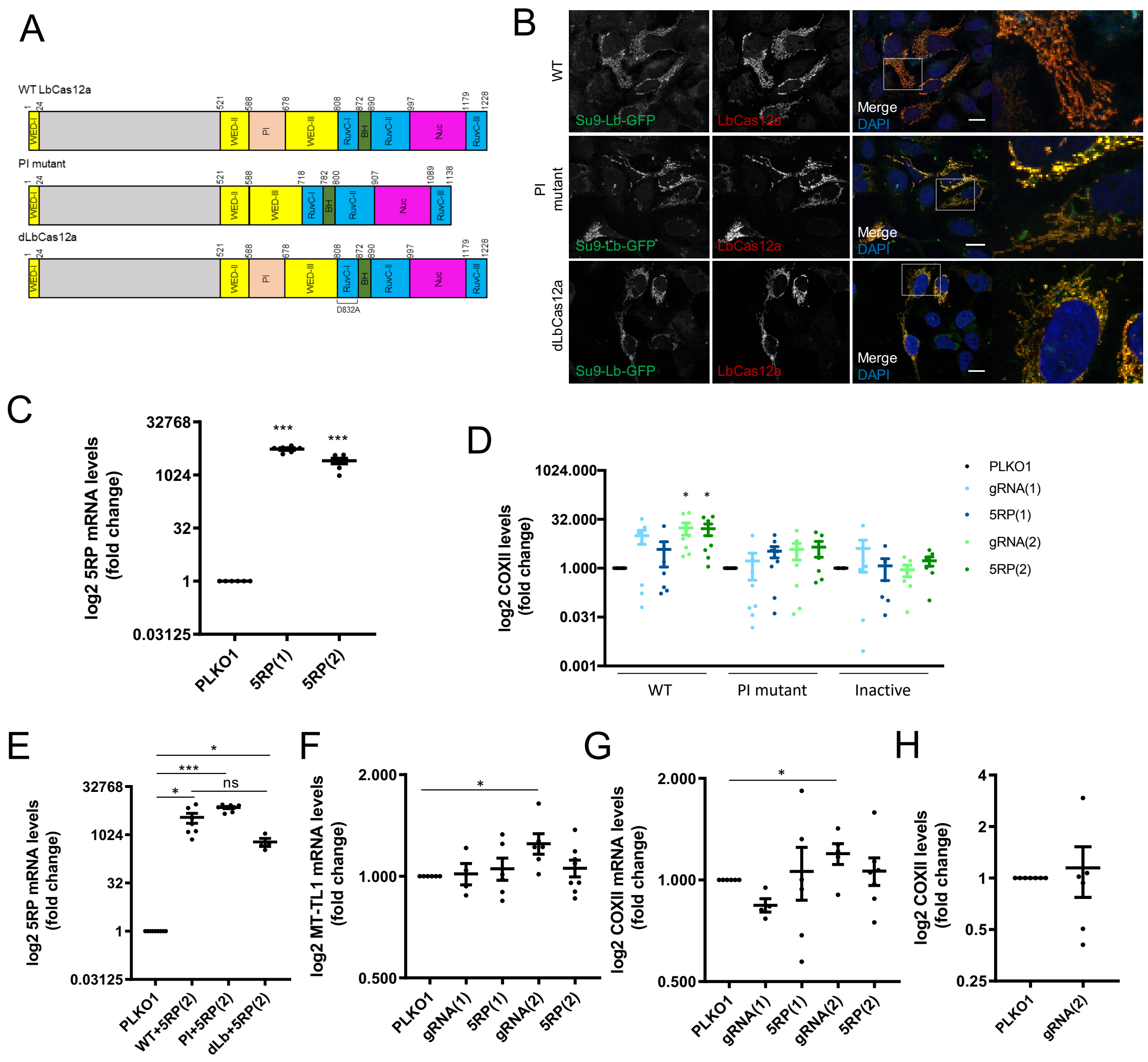

**Figure S8**. Quantitative analysis of mitochondrial DNA and mRNA levels in MELAS cybrids. (A) Schematic diagram of the domain organization of WT LbCas12a, a PI mutant lacking the PAM interacting (PI) domain, and the inactive form dLbCas12a (D832A). (B) Representative images showing expression and mitochondrial localisation of WT, PI mutant and dLbCas12a following co-transfection with PLKO.1 control in MELAS cybrids. (C) qRT-PCR analysis of 5′RP gRNA transcription in MELAS cybrids following PLKO.1, 5′RP(1) or 5′RP(2)/Su9LbCas12a-GFP co-transfection. (D) qPCR analysis of COXII DNA levels following PLKO.1, gRNA(1), 5RP(1), gRNA(2) or 5′RP(2)/WT, PI mutant or dLbCas12a co-transfection. (E-F) qRT-PCR analysis of MT-TL1 (E) and COXII (F) expression in MELAS cybrids following PLKO.1, gRNA(1), 5′RP(1), gRNA(2) or 5RP(2)/Su9LbCas12a-GFP co-transfection. (G) qRT-PCR analysis of 5′RP gRNA transcription in MELAS cybrids following 5′RP(2)/WT, PI mutant or dLbCas12a co-transfection. (H) qPCR analysis of COXII DNA levels following PLKO.1 or gRNA(2)/Su9LbCas12a-GFP co-transfection. Data was normalised to GAPDH. Means ± SEM (n≥6); student's t test: *p<0.05, ** p<0.01 and *** p<0.001 vs. PLKO.1 control.

**Table 1**. Cas9 and Cas12a gRNA/crRNA sequences.

| gRNA/crRNA | Sequence (5′-3′) |
| --- | --- |
| Cas9: | |
| gRNA | GCGCUAAAGAGGAAGAGGACAGUUUUAGAGCUAGAAAUAGCAAGUUAAAAUAAGG CUAGUCCGUUAUCAACUUGAAAAAGUGGCACCGAGUCGGUGCUUUUUU |
| 5' RP | GUCUCCCUGAGCUUCAGGGAGCGCUAAAGAGGAAGAGGACAGUUUUAGAGCUAGA AAUAGCAAGUUAAAAUAAGGCUAGUCCGUUAUCAACUUGAAAAAGUGGCACCGAGU CGGUGCUUUUUU |
| 5'FD (F1D1) | GGCGCAAUCGGUAGCGCUUCGAGCCCCCUACAGGGCUCCACCGCGCUAAAGAGG AAGAGGACAGUUUUAGAGCUAGAAAUAGCAAGUUAAAAUAAGGCUAGUCCGUUAU CAACUUGAAAAAGUGGCACCGAGUCGGUGCUUUUUU |
| 5'AD | GCCUUGUUGGCGCAAUCGGUAGCGCAAAUACAGGGCUCCACGCUAAAGAGGAAGA GGACAGUUUUAGAGCUAGAAAUAGCAAGUUAAAAUAAGGCUAGUCCGUUAUCAAC UUGAAAAAGUGGCACCGAGUCGGUGCUUUUUU |
| Cas12a (1) | AAUUUCUACUAAGUGUAGAUUUAAGAUGGCAGGGCCCGGUAAUC |
| Cas12a (2) | AAUUUCUACUAAGUGUAGAUUGCGAUUACCGGGCCCUGCCAUCU |

**Table 2**. Oligonucleotides for Northern Blotting probes used in **Figure 2**.

| Primer/Oligo | Sequence (5′-3′) |
| --- | --- |
| Cyto $tRNA^{Lys}_{UUU}$ hybridisation probe | ACTTGAACCCTGGACC (16) |
| Mito $tRNA^{Leu}$ hybridisation probe | GAACCTCTGACTCTAAAG (18) |
| Mito $tRNA^{Thr}$ hybridisation probe | TCTCCGGTTTACAAGAC (17) |
| tRK1 hybridisation probe | TGGAGCCCTGTAGGGG (16) |
| gRNA hybridisation probe | GCACCGACTCGGTGCCACTT (20) |

**Table 3**. Parameters that would predict the import efficiency of SpyCas9, SaCas9, LbCas12a, AsCas12a.

| CRISPR Enzyme | Construct | Length / pI signal sequence | Length / pI mature protein | Protein charge | Grand average of hydropathicity (GRAVY) |
| --- | --- | --- | --- | --- | --- |
| **SpyCas9** | COX8a-SpyCas9-GFP | 32 aa / 12.00 | 1616 aa / 8.81 | Positive (+23) | -0.542 |
| **SaCas9** | COX8a-SaCas9-GFP | 32 aa / 12.00 | 1301 aa / 8.98 | Positive (+28) | -0.714 |
| | Su9-SaCas9-GFP | 69 aa / 12.55 | 1297 aa / 9.00 | Positive (+37) | -0.712 |
| | Atg4D-SaCas9-GFP | 41 aa / 11.17 | 1340 aa / 9.13 | Positive (+32) | -0.725 |

| | | | | | |
|---|---|---|---|---|---|
| **LbCas12a** | COX8a-LbCas12a-GFP | 32 aa / 12.00 | 1476 aa / 7.15 | Positive (+4) | -0.564 |
| | **Su9-LbCas12a-GFP** | 69 aa / 12.55 | 1474 aa / 7.33 | Positive (+13) | -0.565 |
| | **Su9-LbCas12a** | | 1263 aa / 8.36 | Positive (+21) | -0.598 |
| | Atg4D-LbCas12a-GFP | 41 aa / 11.17 | 1515 aa / 8.30 | Positive (+8) | -0.574 |
| **AsCas12a** | COX8a-AsCas12a-GFP | 32 aa / 12.00 | 1555 aa / 6.85 | Negative (-1) | -0.472 |
| | Su9-AsCas12a-GFP | 69 aa / 12.55 | 1551 aa / 6.92 | Positive (+8) | -0.476 |
| | Atg4D-AsCas12a-GFP | 41 aa / 11.17 | 1594 aa / 7.94 | Positive (+3) | -0.483 |
